# Task activations produce spurious but systematic inflation of task functional connectivity estimates

**DOI:** 10.1101/292045

**Authors:** Michael W. Cole, Takuya Ito, Douglas Schultz, Ravi Mill, Richard Chen, Carrisa Cocuzza

**Affiliations:** Center for Molecular and Behavioral Neuroscience, Rutgers University, Newark, NJ, 07102, USA; Behavioral and Neural Sciences PhD Program, Rutgers University, Newark, NJ 07102, USA

**Keywords:** functional connectivity, networks, fMRI, computational modeling, method validation

## Abstract

Most neuroscientific studies have focused on task-evoked activations (activity amplitudes at specific brain locations), providing limited insight into the functional relationships between separate brain locations. Task-state functional connectivity (FC) - statistical association between brain activity time series during task performance moves beyond task-evoked activations by quantifying functional interactions during tasks. However, many task-state FC studies do not remove the first-order effect of task-evoked activations prior to estimating task-state FC. It has been argued that this results in the ambiguous inference “likely active or interacting during the task”, rather than the intended inference “likely interacting during the task”. Utilizing a neural mass computational model, we verified that task-evoked activations substantially and inappropriately inflate task-state FC estimates, especially in functional MRI (fMRI) data. Various methods attempting to address this problem have been developed, yet the efficacies of these approaches have not been systematically assessed. We found that most standard approaches for fitting and removing mean task-evoked activations were unable to correct these inflated correlations. In contrast, methods that flexibly fit mean task-evoked response shapes effectively corrected the inflated correlations without reducing effects of interest. Results with empirical fMRI data confirmed the model’s predictions, revealing activation-induced task-state FC inflation for both Pearson correlation and psychophysiological interaction (PPI) approaches. These results demonstrate that removal of mean task-evoked activations using an approach that flexibly models task-evoked response shape is an important preprocessing step for valid estimation of task-state FC.

**Highlights:**

- Computational model shows task inflation of functional connectivity estimates
- Hemodynamic responses cause task activations to further inflate estimates
- Standard approaches to remove task activations leave many false positives
- Methods that flexibly fit hemodynamic response shape effectively correct inflation
- Correction of functional connectivity inflation verified with empirical fMRI data

## INTRODUCTION

Converging evidence across a wide variety of neuroscientific methods applied across multiple species suggests cognition emerges from widespread neural interactions (Cole et al., 2013; Gratton, 2013; Likhtik et al., 2005; M. Siegel et al., 2015). A common way to characterize these cognitive brain network interactions involves estimating task-state functional connectivity (FC) - statistical associations between neural time series during task performance (Friston, 2011; 1994). Typically, such statistical associations are interpreted as evidence of interactions between neural entities (e.g., neurons, local neural populations, brain regions) (M. R. Cohen and Kohn, 2011; Friston, 2011). However, various extraneous variables (e.g., physiological artifacts, in-scanner motion) can confound such inferences (Behzadi et al., 2007; Birn et al., 2006; Power et al., 2012a). We focus here on the possibility that neural activity time-locked to task events (“evoked” activity) is problematic for proper task-state FC inferences.

As Figure 1 illustrates, the proposed issue is that experimenter-controlled task timing creates a temporal pattern in all neural entities active in response to the task, irrespective of whether these neural entities are interacting. This may create statistical associations merely due to similarity with the task timing. For instance, presenting a visual stimulus simultaneously with an auditory stimulus would increase activity simultaneously in primary visual and primary auditory cortices, increasing visual-auditory FC estimates despite no change in interaction between these neural entities. Instead, it would be preferable to remove such task-timing-driven statistical associations, leaving statistical associations to be driven by moment-to-moment (and event-to-event) neural activity fluctuations shared across neural entities (**Figure 1C, 1D**, **& 1E**). Notably, these remaining shared neural activity fluctuations are thought to reflect statistical interactions between the psychological context induced by the task and underlying neural processes (Friston et al., 1997; McLaren et al., 2012), allowing more valid estimation of task-state FC separately from confounding effects of task-evoked activations.

**Figure 1 -.**
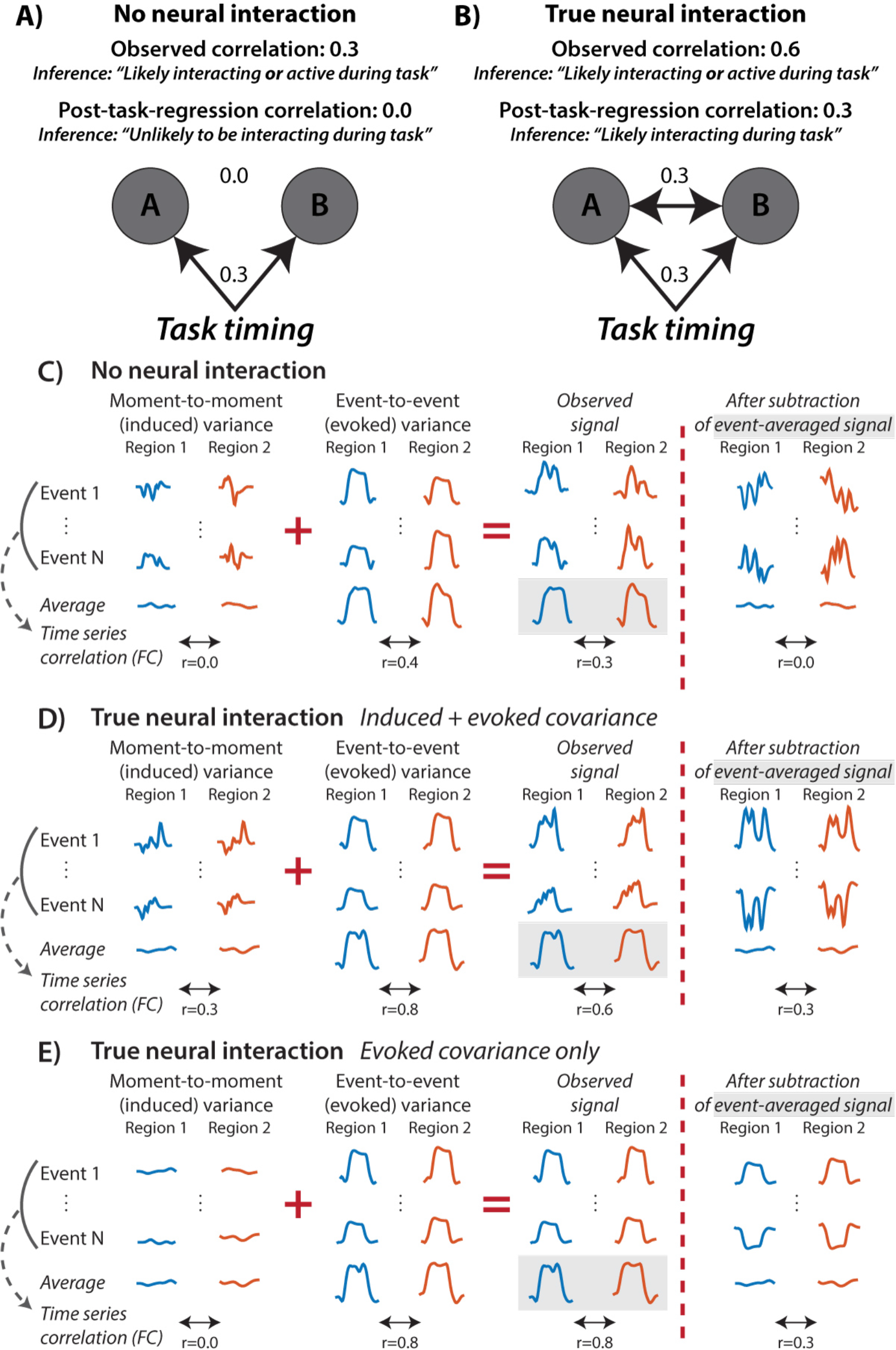
Illustration of the possibility that task-evoked activations are problematic for proper task-state FC inferences. **A**) Graph depicting a scenario with no true neural interaction between A and B. A and B both increase their activity in response to task events, but they do not interact either directly or indirectly. The task event timing nonetheless acts as a confounder to create an artefactual correlation between the neural populations (‘original correlation’). Regressing out the mean task activation (the first-order effect of task) and estimating FC on the residuals (the second-order effect of task; ‘post-task-regression correlation’) removes this artifactual correlation. **B**) Even with a true positive, the task-evoked activity inflates the FC estimate relative to the ground truth interaction. **C**) An illustration (with artificial time series) of signal components underlying the “no true interaction” scenario. “Induced activity” is moment-to-moment variance in brain activity that is not time-locked to task timing. “Evoked activity” is event-to-event (e.g., block-to-block or trial-to-trial) variance in the brain activity that is time-locked to each task event onset. Note that evoked activity always varies in amplitude event-to-event in practice (due to the inherent noisiness of brain processes). Subtracting the mean evoked response from the timeseries before computing the correlation corrects for the inflation. **D**) An illustration of the “true interaction” scenario. **E**) A hypothetical example with only minimal induced variance, illustrating that “true” evoked covariance can drive corrected task-state FC results even after removing mean evoked responses.

A variety of researchers have considered such task-evoked activity to be a confound for FC analyses for both invasive and non-invasive neural recordings, and have implemented various strategies for controlling for this confound during data analysis (Fair et al., 2007; Friston et al., 1997; Gerstein and Perkel, 1972; Kalcher and Pfurtscheller, 1995; Narayanan and Laubach, 2009). However, to our knowledge such task-state FC false positives have not been systematically investigated in either simulations or empirical data. Indeed, many task-state FC studies still do not acknowledge this potential confound. These studies were justified in not worrying about this putative problem, given that it has not been conclusively established in the literature (it has only been *assumed* to be a problem by some researchers). Thus, while this problem may already seem very real to some researchers, many remain skeptical as to its existence. Moreover, beyond establishing the central problem of task FC inflation (and its extent), there is a need to systematically evaluate methods of correction, given the lack of methodological consensus among task-state FC studies that have acknowledged the problem.

Here we sought to more conclusively test the hypothesis that the task-timing confound exists for task-state FC (for all neural recording methods, not just functional MRI (fMRI)), using computational modeling. Additionally, we tested the hypothesis that the task-timing confound is more pronounced for fMRI data. This hypothesis is based on the temporally-extended nature of hemodynamic response function (HRF) profiles and their similarity across neural entities (despite not being identical (Handwerker et al., 2004)). These HRF features may increase the statistical similarity among task-timing-locked evoked fMRI responses. We reasoned that strategies to correct for the task-timing confound would be even more critical for fMRI than for other methods if this feature of fMRI data inflates the confounding effect of task timing.

As a starting point, we focused primarily on block (rather than event-related) cognitive paradigms. This was done for both theoretical and practical reasons. Theoretically, neural time series during block designs are less likely to be strongly influenced by task timing effects, due to fewer rest-to-task state transitions (Gitelman et al., 2003; O’Reilly et al., 2012). This makes block designs a stronger test of the generalizability of our task FC inflation hypothesis. In other words, if a task-timing confound is found with block designs it is very likely to also be a problem for event-related designs. Practically, the sluggishness of fMRI hemodynamics makes it difficult to separate temporally proximal events in event-related designs, making it challenging to obtain clean estimates of task-state FC for each task condition. This reflected our approach of isolating task epochs from other rest or task epochs for all analyses, which reduces (but does not eliminate, due to temporal autocorrelations) the chance of state transitions driving observed false positives. It will be important for future studies to verify that the conclusions drawn here generalize to event-related designs.

The standard approach to correct for the task-timing confound is to fit an event-averaged general linear model (GLM) of the task events either simultaneously with task-state FC (as with psychophysiological interaction; PPI) (McLaren et al., 2012; O’Reilly et al., 2012) or calculate FC estimates using the residual time series of such a model (Al-Aidroos et al., 2012; Cole et al., 2013; Gratton et al., 2016; Summerfield et al., 2006). To clarify the rationale behind these approaches (following the parlance of electroencephalography), inflation of task FC by task activations is corrected by removing the mean “evoked” responses (time-locked to task events) so as to isolate “induced” responses (responses influenced by task events but that vary in timing across multiple instances of those events) (Tallon-Baudry and Bertrand, 1999). Note that evoked responses that vary in amplitude across events remain in addition to induced responses (Truccolo et al., 2002a) (**Figure 1E**). Neural time series correlations that remain after removing the cross-event mean response are termed (perhaps inappropriately) “noise correlations” in the non-human animal neurophysiology literature (Cafaro and Rieke, 2010; M. R. Cohen and Kohn, 2011). One goal of the present study is to determine whether only removing cross-event mean (evoked) responses is adequate for eliminating task-activation-driven FC inflation. A second goal is to evaluate the effect of different methods of modeling the HRF on the reduction of task-inflation of FC estimates.

An overview of our approach is outlined in **Table 1**. We began by testing for the existence of the proposed task-state FC confound using a highly simplified simulation. This was followed by a more complex simulation utilizing a neural mass computational model that included more features of real neural data. Once the task-state FC confound was identified in simulated fMRI (and non-fMRI) data, we tested a variety of methods to correct for the confound. Once a confound-correction method was identified, we tested its efficacy in real fMRI data. Critically, demonstrating that this confound-correcting method has a strong impact on results with real fMRI data would provide more conclusive evidence that the confound exists and that correcting it matters in practice.

**Table 1 -.** Overview of study objectives and approaches. All three approaches were used to assess the two study objectives. However, lack of “ground truth” (the true FC values independent of noise and artifacts) with empirical fMRI data limits the ability to verify whether task activation increases FC false positives. In contrast, the two computational models provided “ground truth” scenarios for verifying activation-induced FC inflation. Note that in the absence of false negatives driven by FC inflation correction methods (as demonstrated in the neural mass model) one can use the reduction in empirical task-state FC estimates as indirect evidence of FC inflation.

| **Approach** | **Objectives** |  | **Advantages** | **Disadvantages** |
| --- | --- | --- | --- | --- |
|  | *Determine whether task activations increase FC false positives* | *Test methods to reduce activation-induced FC inflation* |  |  |
| 1) Minimal model | ✓ | ✓ | Minimal assumptions; know "ground truth" | Missing key features of real FC |
| 2) Neural mass model | ✓ | ✓ | Includes key features of real FC; know "ground truth" | More assumptions (than minimal model & empirical data) |
| 3) Empirical fMRI data | ~ | ✓ | No assumptions (actual data of interest) | Do not know "ground truth" (though "ground null" estimated; Figure S2) |

## METHODS

### Minimal model

We began with a very simple test of the hypothesized task-timing-induced FC inflation effect. This involved creating two Gaussian random time series (mean = 0, standard deviation = 1) with very low correlation (r=-0.10), followed by adding a value of 1.0 during two “task” blocks. This can be thought of as an increase in activity/excitability for both “nodes”. Task blocks lasted 30 time points each, with 30 time points of “rest” before and after. The model and analysis of model data were implemented in Python (version 2.7.13) with modules numpy (version 1.12.1) and matplotlib (version 2.0.0). HRF convolution was performed using a standard double-gamma HRF shape, treating each time point as representing 1 second. Pearson correlation (numpy function corrcoef) was used to estimate FC between the time series. Code to implement the model and run all analyses are available here: https://github.com/ColeLab/TaskFCRemoveMeanActivity/blob/master/minimalmodel/MinimalModel.ipynb

### Neural mass model

To extend the findings based on the minimal model, we developed a neural mass model to simulate the large-scale activity and interaction patterns of sets of thousands of neurons, based on standard neural mass models (Cole et al., 2016a; Hopfield, 1984; Ito et al., 2017; Wilson and Cowan, 1972). We sought to optimize the model simultaneously for simplicity and biological interpretability. We expected simplicity to increase the interpretability of results and computational tractability, while we expected biological interpretability to facilitate the relationship between the simulation results to neuroimaging results. The core of the model is a standard firing rate model, which uses a sigmoid transfer (input-output) function (Cole et al., 2016a; Hopfield, 1984; Ito et al., 2017; Wilson and Cowan, 1972). Using a standard firing rate model increased the simplicity of the model compared to some alternatives, while remaining biologically plausible based on evidence that neural populations exhibit a sigmoid-like transfer function reflecting variability in the exact firing threshold across individual neurons (Hopfield, 1984).

We defined each node’s output as:

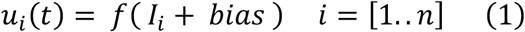

where *u_i_* is the output activity (population spike rate) for unit *i* at time *t*, *I*_*t*_ is the input (population field potential) as defined below, and *bias* is the bias (population resting potential, or excitability).

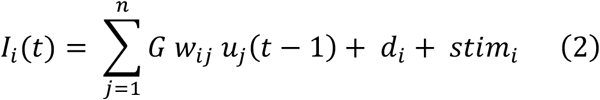

where *I*_*i*_(*t*) is the input (population field potential) for unit *i* at time *t, G* is the global coupling parameter (a scalar influencing all connection strengths), *w_ij_* is the synaptic weight from unit *j* to *i*, *u*_j_(*t* − *τ_ij_*) is the output activity from unit *j* at the previous time step (*τ_ij_* was set to 1 for simplicity), *d*_*i*_ is spontaneous activity (independent Gaussian random values across nodes), and *stim*_*i*_ is task stimulation (if any).

The initial condition (input at time point 0) of each unit for each simulation was set to a Gaussian random value (mean 0, standard deviation 1).

The sigmoid *f*(*x*) (population threshold) in the node output equation above is defined as:

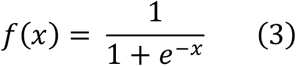

We reduced the arbitrariness of model parameter selection using a principled parameter search. Parameters for the model were determined based on optimizing for high task-state FC relative to resting-state FC (without fMRI simulation). Specifically, we selected parameters that generated the highest average task-state FC (relative to the average resting-state FC) among the first 50 nodes in the 300-node network described in the next section. Optimizing for only a portion of the entire network reduced the chance that the optimization overfit to the particular network structure. Rerunning the model with multiple initial random conditions (for the main analyses) also ensured overfitting was not an issue. Notably, we did not optimize for higher task-state FC false positives nor for high fMRI-based FC, such that we could test for fMRI-based false positives as a hypothesis independent of how the model parameters were chosen. The parameter search involved all permutations of the model parameters varied, with the following ranges: *G*=1 to 10, self-connections (diagonals in *w*)=0 to 10, *bias*=-15 to 0, *d*=1 to 10, *stim*=0.1 to 1.0 (in 0.1 increments).

Settings used for the model: *d_i_* was a Gaussian random value with mean 0 and standard deviation 3, *G* was set to 5, *bias* was set to -5, *stim* was set to 0.3, and all self-connections (diagonals in *w*) were set to 1. Setting the self-connection above 0 reflects the theoretical neurons within the modeled neural population having synaptic connections among each other, such that the same outputs sent to other units also affect the unit that sent it. The model was implemented in Python (version 2.7.13).

### The model’s network organization

The model’s network included 300 nodes, selected to be in the same range as some recent estimates of the number of functional regions in human neocortex (Glasser et al., 2016; Van Essen et al., 2012). This 300-node network was given a functional network community structure, based on empirical evidence of such large-scale network structure in the human brain (Ito et al., 2017; Power et al., 2011; Spronk et al., 2017). Briefly, the construction and running of each “subject’s” network went as follows: 1) Build structural and synaptic connectivity network architecture, 2) Simulate neural activity during both a resting-state run and a task-state run (detailed below), 3) Simulate fMRI data collection by converting each node’s “input” time series to fMRI via convolving with an HRF and downsampling the resulting time series.

Network construction involved a series of steps (Cole et al., 2016a), with the construction of the network model randomly initialized separately for each “subject”. First, there was a 10% probability of any node in the network connecting to any other. Next, three structural communities were created by increasing the probability of connectivity within each set of 100 nodes to 50%. This was then converted to a synaptic connectivity matrix by adding a Gaussian random value to each structural connection (mean of 1, standard deviation of 0.001). The first structural community was then split into two “functional” communities (defined based on synaptic weights, rather than the mere existence of a connection) by multiplying the synaptic weights among the first 50 nodes (and, separately, the second 50 nodes) by 1.2 and multiplying the connections to/from the first and second 50 nodes by −0.2. Next, all connections to/from the final 100 nodes and all other nodes were multiplied by 0, completely isolating the final community from the rest of the network. Finally, each node’s synaptic connectivity was normalized such that all inputs summed to 1.0. Input weight normalization is thought to be a biologically realistic process (e.g., via each neuron regulating the number of channels at each synapse) (Barral and D Reyes, 2016).

Task stimulation amplitude targeted 25 nodes in the first and last network communities. Note that the setting of the bias to -5 was consistent with units starting out at a near-0 firing rate (given the sigmoid activation function that was used), modeling most neurons within a modeled population being at a sub-threshold resting potential. Model conversion to fMRI data involved convolution of variable HRFs with the input time series from each node. The HRF differed for each simulated subject and each region, though it differed more between subjects than between regions, consistent with empirical evidence (Handwerker et al., 2004). A standard double-gamma HRF function was used in all cases, with variation in the double-gamma parameters across nodes and subjects. Specifically, the values for peak time (3 to 9 in increments of 0.5 s), undershoot time (3 to 17 in increments of 0.5 s), and undershoot ratio (0 to 1 in increments of 0.1) of a double-gamma HRF were varied randomly (uniform distribution) by subject. Then, each node had these three parameters varied from a given subject’s selected values based on a Gaussian random distribution centered on 0 with a standard deviation of 1, with that value being the array index selecting from the set of allowed values for each parameter (as indicated in the previous sentence). Note that results were similar without HRF variability (i.e., with the same non-canonical HRF shape used for all subjects and all regions). HRF convolution was followed by sampling (selecting a single time point) of the convolved time series at a time to repetition (TR) of 0.785 seconds, in the range of multiband fMRI protocols (Chen et al., 2015).

The model was implemented with 24600 time steps per “run”, with each time step conceptualized as 50 ms, such that the total simulated time was conceptualized as 20.5 minutes in duration. Each run was implemented across 24 “subjects”, with a separate random seed used for each subject for the spontaneous activity. The first run consisted of a resting-state simulation with no task stimulation. The second run consisted of a task-state simulation, with 6 task “blocks” of 2.5 minutes of constant stimulation of the two sets of nodes indicated above. There was 30 seconds of non-stimulation before and after each task block. All FC analyses used the time points included in the 6 task blocks, ignoring the inter-block periods.

### FC estimation

Estimates of time series association were calculated using either MATLAB (version R2014b) or R (version 2.15.1). Pearson correlation was calculated as:

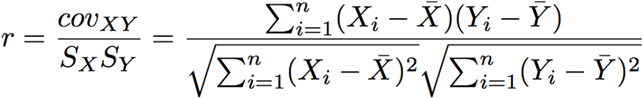

Where *S* is the time series standard deviation, *cov* is the time series covariance, *X* and *Y* are brain activity time series, *n* is the number of time points, and *X̄* and *Ȳ* are the time series means. Most analyses also involved the Fisher’s z-transform of the resulting Pearson correlation. The Fisher’s z-transform:

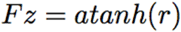

Psycho-physiological interaction (PPI) was estimated using simple linear regression, which is equivalent to:

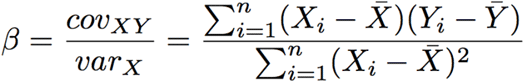

Where *var* is the time series variance. The beta for each condition was estimated separately for each condition, consistent with generalized PPI (McLaren et al., 2012).

### Empirical fMRI data collection

The empirical fMRI dataset was collected as part of the Washington University-Minnesota Consortium Human Connectome Project (HCP) (Van Essen et al., 2013). These data are publicly available, accessible at https://www.humanconnectome.org. Participants were recruited from Washington University (St. Louis, MO) and the surrounding area. All participants gave informed consent. The data used were selected by the HCP as the “100 unrelated” dataset, consisting of data from 100 participants with no family relations. Data from 25 subjects were not used because of excessive in-scanner movement (defined as over 50% of volumes in any run with mean framewise displacement > 0.25 mm) for these subjects, such that data from 75 subjects were included in the final analyses. Framewise displacement was calculated as described by Power et al. (2012b), with a low-pass filter of 0.3 Hz applied as suggested by Siegel et al. (2016) for multiband fMRI data.

Whole-brain echo-planar imaging acquisitions were acquired with a 32 channel head coil on a modified 3T Siemens Skyra with TR = 720 ms, TE = 33.1 ms, flip angle = 52°, BW = 2290 Hz/Px, in-plane FOV = 208 × 180 mm, 72 slices, 2.0 mm isotropic voxels, with a multi-band acceleration factor of 8 (Ugurbil et al., 2013). Data were collected over two days. On each day 28 minutes of rest (eyes open with fixation) fMRI data across two runs were collected (56 minutes total), followed by 30 minutes of task fMRI data collection (60 minutes total). Each of the 7 tasks was completed over two consecutive fMRI runs. Resting-state data collection details for this dataset can be found elsewhere (Smith et al., 2013), as can task data details (Barch et al., 2013).

This dataset included 24 task conditions across seven tasks, with each task completed by all subjects. All seven tasks were block designs, with varying block durations and delays across the tasks. Task timing details, as described by (2013), are included in **Table 2**.

**Table 2 -.**
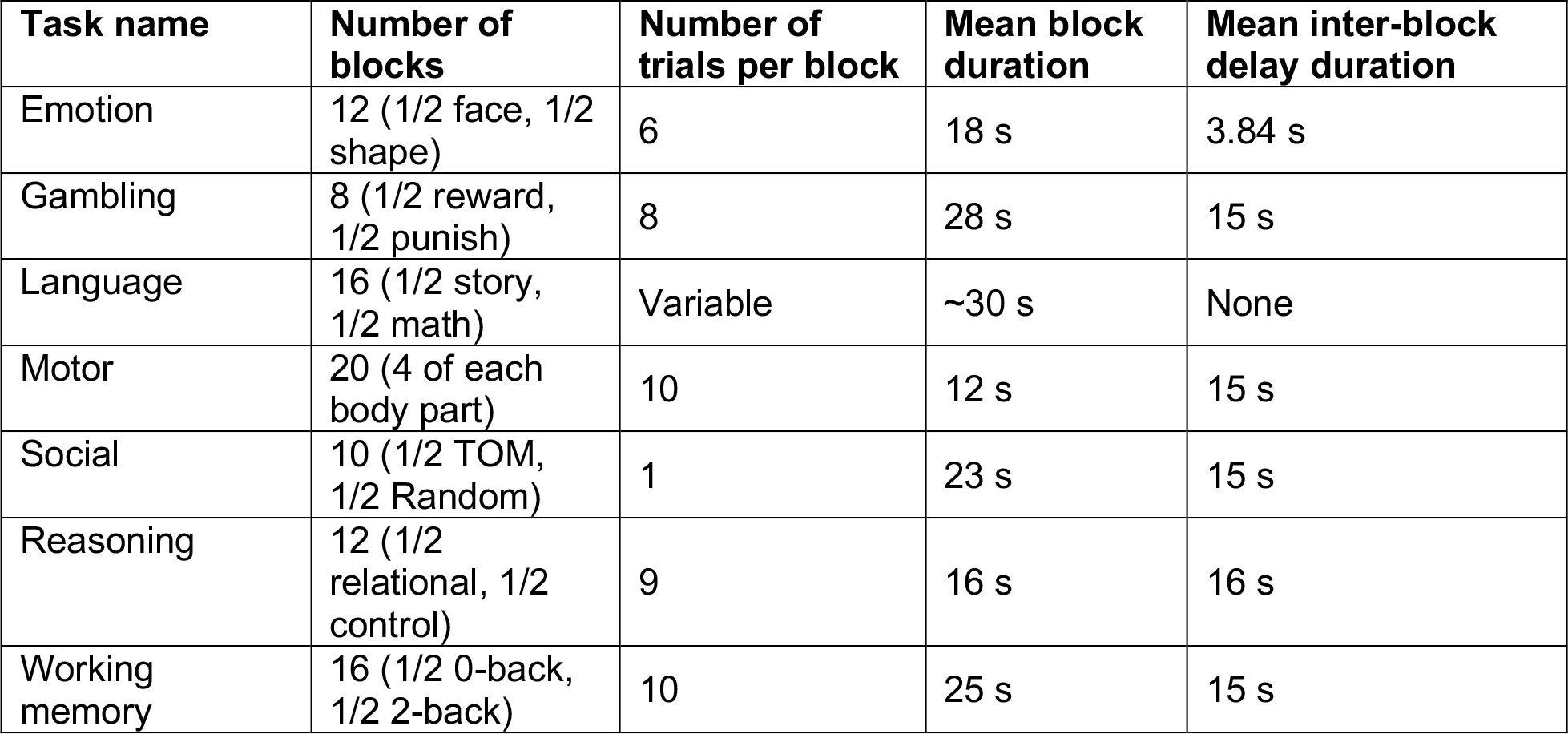
Empirical fMRI task timing details. See (Barch et al., 2013) for more information regarding task timing in the Human Connectome Project dataset.

| Task name | Number of blocks | Number of trials per block | Mean block duration | Mean inter-block delay duration |
| --- | --- | --- | --- | --- |
| Emotion | 12 (1/2 face, 1/2 shape) | 6 | 18 s | 3.84 s |
| Gambling | 8 (1/2 reward, 1/2 punish) | 8 | 28 s | 15 s |
| Language | 16 (1/2 story, 1/2 math) | Variable | ~30 s | None |
| Motor | 20 (4 of each body part) | 10 | 12 s | 15 s |
| Social | 10 (1/2 TOM, 1/2 Random) | 1 | 23 s | 15 s |
| Reasoning | 12 (1/2 relational, 1/2 control) | 9 | 16 s | 16 s |
| Working memory | 16 (1/2 0-back, 1/2 2-back) | 10 | 25 s | 15 s |

### Empirical fMRI dataset analysis

The empirical dataset preprocessing consisted of standard functional connectivity preprocessing (typically performed with resting-state fMRI data), with several modifications given that analyses were also performed on task-state data. Resting-state and task-state data were preprocessed identically to facilitate comparisons between them. Spatial normalization to a template (MSM-sulc), motion correction, intensity normalization (normalized to a 4D whole brain mean of 10,000) were already implemented in a minimally-processed version of the empirical fMRI dataset described elsewhere (Glasser et al., 2013), so we began preprocessing with this version of the data. With the surface (rather than the volume) version of the minimally preprocessed data, we used custom scripts in MATLAB to additionally remove nuisance time series (motion, ventricle, and white matter signals, along with their derivatives) using linear regression, and remove the linear trend for each run. A low-pass temporal filter was not applied due to the possible presence of task signals at higher frequencies (e.g., relative to slow resting-state fluctuations).

Data were sampled from a set of 360 brain regions (rather than individual voxels/vertices) to make inferences at the region and systems levels. We used an independently-identified set of putative functional brain regions (Glasser et al., 2016) so as to reduce any potential circularity in analyses (Kriegeskorte et al., 2009). The use of this parcellation also reduces the chance of combining signals from multiple functional regions as compared to anatomically-defined parcellations (Wig et al., 2011). These brain regions were identified using parcellation of a variety of data types, including resting-state functional connectivity, task activation, and myelin maps (Glasser et al., 2016). Data were summarized for each region by averaging signal in all vertices falling inside each region.

Preprocessing was carried out using Freesurfer (version 5.3.0-HCP), FSL (version 5.0.8), and custom code in MATLAB 2014b (Mathworks) for the 7-task dataset (using the minimally preprocessed version of the data (Glasser et al., 2013)). Further analysis was carried out with MATLAB and R.

We estimated FC using Pearson correlations and regressions between time series from all pairs of brain regions using MATLAB (version R2014b). For Pearson correlations, all computations used Fisher’s z-transformed values. FC estimation was straightforward for resting-state data, as there were no additional steps after preprocessing prior to calculating these values. For task data there were additional steps related to task activation regression, as described in the following section.

FC differences were assessed using two-way t-tests paired by subject. Multiple comparisons were corrected for using false discovery rate (FDR) (Genovese et al., 2002). When comparing task-state FC to resting-state FC estimates the number of time points contributing to those estimates were matched. The beginning of the first resting-state fMRI run was used in all cases, due to the increased likelihood of subjects falling asleep later in the rest run (Tagliazucchi and Laufs, 2014).

### Task-activation regression for task-state FC

Cross-event (trial and block) mean responses during task fMRI might unduly influence task-related changes in FC. This was rigorously tested using computational modeling, which informed our empirical fMRI data analysis. We sought to suppress or remove such influences with task regression techniques. This involved running standard fMRI general linear model (GLM) analysis, and calculating FC based on the residuals. Specifically, each region’s task time series was modeled using a GLM, with a distinct model depending on the analysis (as described below). To improve removal of task-related activation variance, a separate regressor was included for each task condition (e.g., face stimuli vs. tool stimuli in the N-back task; 24 task conditions total). Note that regressing out task events using GLM removes the cross-event response means, retaining event-to-event and sub-event fluctuations in time series such that these sources of variability likely contribute the most to task FC estimates (Rissman et al., 2004a; Truccolo et al., 2002b). The residuals from this regression model were used for FC estimation, restricted to time points corresponding to the current task. A standard hemodynamic lag was included when determining task timing, by convolving the timing with a canonical HRF and selecting time points with a value above 0.

FC estimation was conducted along with no task regression, canonical HRF task regression, constrained basis set task regression, or finite impulse response (FIR) task regression. Other than the task regression step, all steps were identical in the no-task-regression case as when task regression was used. Canonical HRF task regression involved use of the SPM software function spm_hrf.m with the default parameters to create the HRF. This HRF was then convolved with each of the 24 task condition time series, then fit using ordinary least squares regression in MATLAB (function regress.m). Constrained basis set task regression involved creation of a set number of basis set regressors (either 5 or 28) in the FLOBS interface in FSL software (version 5.0.8; default parameters) (Woolrich et al., 2004). Note that the first three basis function regressors are highly similar to the canonical, time, and dispersion derivatives often used together to model task activations in SPM software (Woolrich et al., 2004). These basis set functions were then convolved with each of the 24 task condition time series before fitting them to the brain region time series (identically to the canonical HRF approach).

The FIR task regression approach involved fitting the cross-trial/cross-block mean response for each time point in a set window length that is time-locked to the trial/block onset for a given task condition. This allows the fit to be completely flexible with regard to the HRF response shape, so long as it is consistent across trials/blocks for that condition. Each of the 24 task conditions were fit with a series of regressors, one per time point. Each condition’s window length matched the duration of the events, with an additional 18 s (25 regressors) added to account for the likely duration of the HRF. Note that FIR regression is nearly identical to simply subtracting the mean evoked response (see Figure 1), which is a standard method in the spike correlation literature for removing task-evoked activation-driven inflations (Cafaro and Rieke, 2010). The primary difference is that FIR regression can better deal with overlap in observed task events (Miezin et al., 2000), which is especially useful for fMRI data given the sluggishness of hemodynamic responses.

## RESULTS

### A minimal model to demonstrate activation-induced false positives in task-state FC estimates

We sought to determine the efficacy of standard task-state FC estimation methods, specifically the possibility of FC false positives arising from task-evoked activations (mentioned as a likely possibility in many previous studies). We were interested both in the activation-induced inflation effect generally (regardless of data collection method), as well as any fMRI-specific effects. We began by using an extremely minimal test of the proposed FC-estimate inflation, in the hopes of establishing the effect theoretically and identifying likely causes of the inflation.

The easy-to-read Python code used for the minimal model (including figure creation) can be accessed here in a Jupyter Notebook: https://github.com/ColeLab/TaskFCRemoveMeanActivity/blob/master/minimalmodel/MinimalModel.ipynb. This minimal demonstration involved creating two Gaussian random time series with very low correlation (r=-0.10) (**Figure 2A**), followed by adding a value of 1.0 for two “task blocks” for both time series (**Figure 2B**). This can be thought of as an increase in activity for both “nodes”. Consistent with the hypothesized task-timing confound, the time series went from an original whole-time-series correlation of r=−0.10 to a whole-time-series correlation (i.e., not restricted to just the “task” portion of the time series) of r=0.79. This increase held despite the absence of a correlation in the “task” segment (r=−0.07) (**Figure 2C**). This demonstrates the importance of isolating task from non-task time periods when calculating task-state FC, since transient rest-to-task activity transitions can drive overall correlation increases. In more realistic neural time series (see the next section), however, temporal autocorrelation would prevent such clean separation of task and non-task time periods. Thus, both fMRI and non-fMRI data likely suffer from task-state FC inflation due to timeseries autocorrelation induced by the onset of task events (Truccolo et al., 2009) (we explore this possibility in the subsequent section).

**Figure 2 -.**
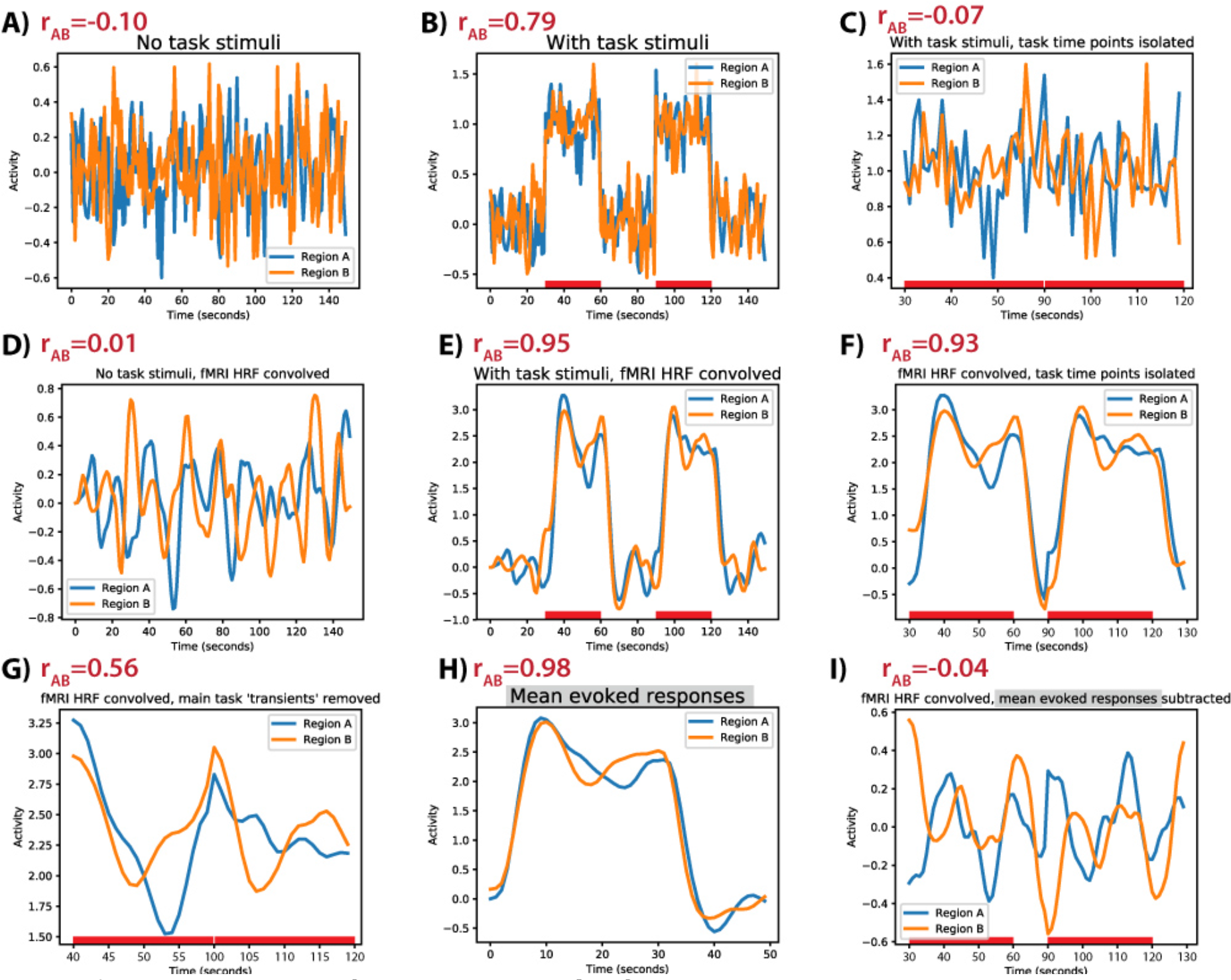
Minimal model: fMRI task-state FC inflation is primarily driven by HRF convolution (temporal autocorrelation), and inflation is corrected by subtraction of mean evoked responses. **A**)Two Gaussian random time series were generated to simulate spontaneous activity in two neural regions that are not “truly” interacting at the neural level. Their correlation is shown in the upper-left corner (as in all other panels). **B**) A “task” was simulated by adding activity in two task blocks. This increased the interregion correlation substantially, indicating the critical role of rest-to-task state transitions in driving correlations. **C**) Simply isolating the task time points (removing the rest-to-task state transition) removed the correlation inflation. **D**) The identical time series in panel A convolved with a standard HRF to simulate the fMRI BOLD response. **E**) An HRF-convolved version of the time series in panel B. **F**) Unlike the “neural” time series, isolating task time points in the “fMRI” time series did not remove the correlation inflation. **G**) Removing the block start and stop transients reduced the correlation inflation, but it was still substantially inflated. **H**) The mean evoked response for each region. **I**) Subtracting the mean evoked response from each region completely removed the correlation inflation in the “fMRI” data.

We next investigated the effect of fMRI (hemodynamics) on task-state FC estimates. This involved convolving the exact same time series with a canonical HRF. Convolution did not substantially change the non-“task” correlation (r=0.01) (**Figure 2D**), but the “task” correlation was highly inflated (r=0.95) (**Figure 2E**). Unlike the “neural” time series (**Figure 2C**), this inflation remained even after isolating the task time points (as well as 10 time points to account for HRF lag) (**Figure 2F**). This demonstrates that the correlation inflation is strongly driven by an interaction between an increase in time series amplitude and HRF convolution.

We next sought to test if subtracting the mean evoked responses (**Figure 2H**) from each task event (equivalent to task GLM regression) would reduce the FC inflation. This is the standard approach to reduce potential activation-induced inflation of task FC estimates in the fMRI (Cole et al., 2013; Gratton et al., 2016) and spike correlation literatures (Averbeck et al., 2006; Grün, 2009). This involved simply averaging the time-locked activity across task blocks and subtracting the mean evoked time series from each task block. As expected, the “task” correlation was substantially reduced (r=−0.04) **(Figure 2I**). This demonstrates the efficacy of task regression (mean evoked response subtraction) for reducing task-amplitude-induced correlation/FC inflation.

### Neural mass model: Testing for false positives

While the simplicity of the prior demonstration gave it clarity, it lacked many features of real neural interactions. Therefore, we next characterized the task-timing-induced false positives using a more realistic neural mass computational model with biologically-interpretable parameters. This involved a standard neural mass model (Cole et al., 2016a; Hopfield, 1984; Ito et al., 2017). Unlike the minimal model in the previous section, the neural mass model provides: interactions among neural units (allowing us to test for false negatives), enough neural masses to plausibly match the number of functional cortical regions in humans (Glasser et al., 2016), large-scale network structure (Power et al., 2011; Yeo et al., 2011), hemodynamic variability (Handwerker et al., 2004), event-to-event variation in neural signals (in addition to moment-to-moment variation) (Fox et al., 2007; 2006), and temporal autocorrelation (Murray et al., 2014; Truccolo et al., 2009).

We constructed a series of large-scale network communities, given the presence of such communities in many real-world networks (Girvan and Newman, 2002) including the human brain (Power et al., 2011; Yeo et al., 2011). We began by making three structural communities of 100 nodes each (**Figure 3A**; see Methods) (Cole et al., 2016a). Importantly, we removed all structural connections to/from the last community (the “no connectivity zone”), allowing us to test for false positives in subsequent analyses (see upper-right corner of **Figure 3A**).

**Figure 3 -.**
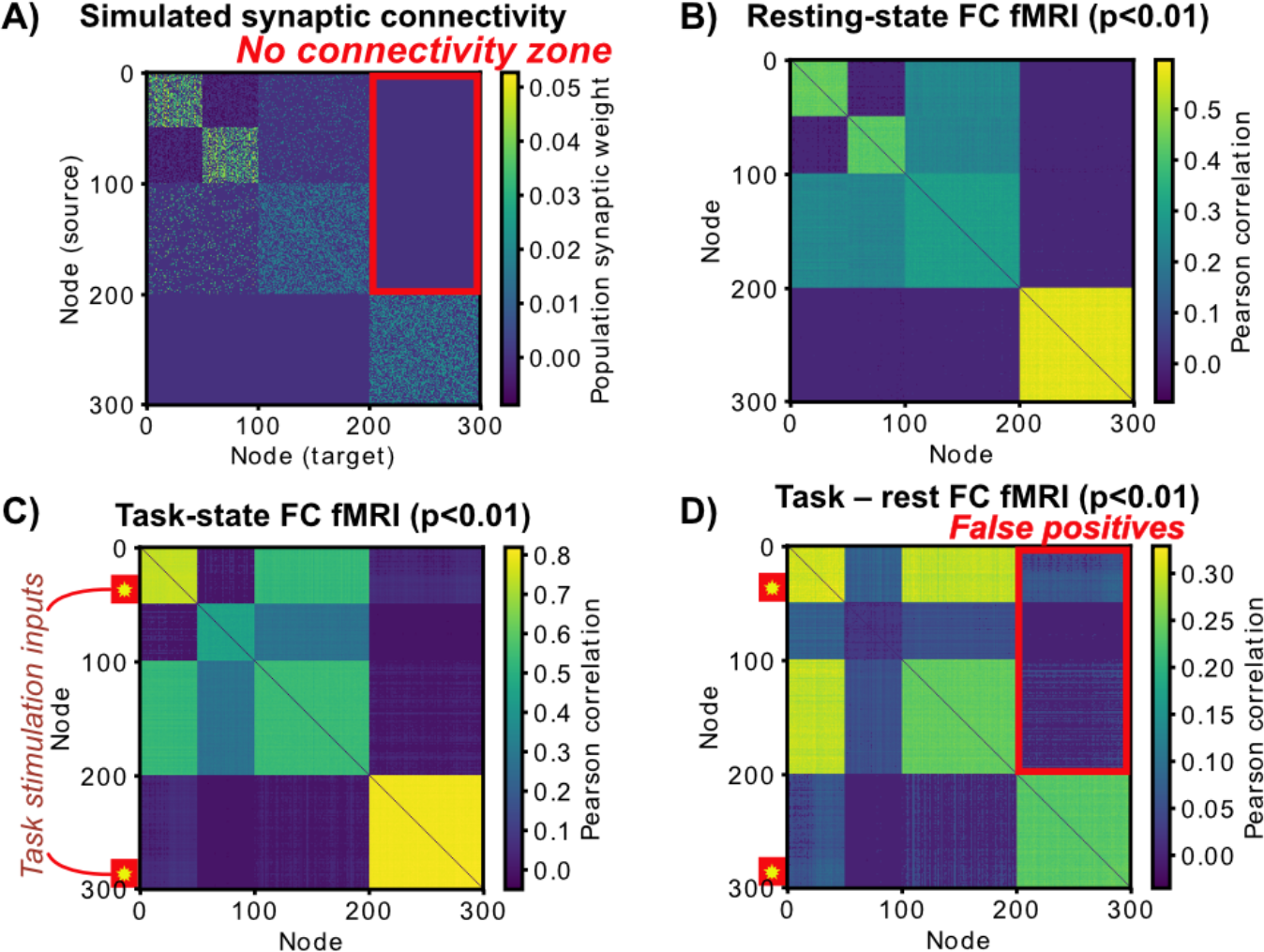
The neural mass model, with fMRI simulation and “no connectivity zone” to test for false positives. **A**) Three structural communities were constructed (100 nodes each), with the first community split into two communities via synaptic connectivity. The first and second structural communities had random connectivity (10% density), while the third community had no connections with the rest of the network. Connections to/from the third community acted as tests for false positives in subsequent simulations (the “no connectivity zone“). **B**) We simulated fMRI by convolving the input time series of each unit with a hemodynamic response function (HRF) and downsampling (every 785 ms). Spontaneous activity without task stimulation was used to produce this FC matrix. T-tests vs. 0 were based on across-subject variance, with each “subject” being a random initialization of the synaptic connectivity matrix and spontaneous activity. Note the low false positive rate (0.81%) (i.e. the lack of significant connections showing up in the ground truth “no connectivity zone”). **C**) Two populations of 25 nodes (indicated by yellow stars) were stimulated simultaneously across 6 task blocks. Two completely unconnected communities were stimulated to test for false positives. Note the increase in false positive connections in the “no connectivity zone” (41%). **D**) T-tests indicated an inflated false positive rate of 40% when comparing task FC to rest FC. Note that without fMRI simulation (i.e., no HRF or downsampling) the false positive rate was 1.99%.

We next used the neural mass model to simulate collection and analysis of resting-state FC with fMRI (**Figure 3B**). To simulate fMRI data, the input (population field potential) time series convolved with an HRF and down sampled. We found that the resting-state FC matrix was significantly similar to the large-scale structure of the synaptic connectivity matrix (mean r=0.47, t(23)=266, p<0.00001). Further, there were minimal false positives (0.8%) in the “no connectivity” zone at a t-test threshold of p<0.01. As is standard for tests for false positives when the null is known to be true, correction for multiple comparisons was not applied, since it would complicate calculation of the false positive rate. Given that we use p<0.01, one can interpret any false positives beyond the 1% rate as true false positives.

We then simulated task-state FC by stimulating two sets of units, in the first and last functional communities (**Figure 3C**). Task stimulation consisted of a small constant input (0.3) across 50 nodes (25 for each of the two communities) in six task blocks. Only the on-stimulation times were analyzed for task FC to reduce the influence of on/off task transients. Increased task-state FC compared to resting-state FC was widespread (**Figure 3D**). This is consistent with the observation of task-state FC across a wide variety of brain systems and tasks in the fMRI literature (Cole et al., 2014; Krienen et al., 2014). However, it was apparent that a large number of false positives were present in the “no connectivity” zone: 42.58% false positives for task vs. rest FC (p<0.01). This appeared to be driven primarily (but not exclusively) by the fMRI simulation, since the false positive rate was only 1.99% (task vs. rest, p<0.01) with the same data prior to fMRI simulation.

### Neural mass model: Testing for correction of the false positive rate

Given verification of a systematic inflation of the false positive rate we next tested proposed approaches to correcting this false positive rate. These typically involve regressing out the task timing, which involves using the residuals of a GLM as the time series to compute task-state FC. This is very similar to simultaneous fitting of task-state FC and task activations when using PPI (Friston et al., 1997; McLaren et al., 2012; O’Reilly et al., 2012). When an finite impulse response (FIR) GLM is used (Cordova et al., 2016; Fair et al., 2007; Norman-Haignere et al., 2012), this is also very similar to simply subtracting the mean evoked response in the spike correlation literature (Averbeck et al., 2006; Grün, 2009). Critically, however, since our simulations provided “ground truth” knowledge of the false positive rate we were able to validate the approaches and verify their efficacy for reducing the false positive rate.

We began by using the most common approach for reducing false positives - fitting the “canonical” HRF shape to remove cross-event mean response correlated with task timing (**Figure 4A**). This is the same HRF shape used in PPI (O’Reilly et al., 2012) and related approaches (Cole et al., 2014). We found that task regression with this canonical HRF shape reduced the false positive rate somewhat but failed to bring it below the 1% specified by the p-value threshold (p<0.01): 20.34% false positive rate (**Figure 4B**). We next used a worst-case scenario “flipped” HRF shape to determine if having an approximately-correct HRF shape (as with the canonical HRF) mattered for reducing the false positive rate (**Figure 4A**). We found that using the wrong HRF shape did a worse job of reducing the false positive rate than the canonical HRF (**Figure 4C**): 25.43% false positive rate. This suggests that the relative accuracy of the HRF shape matters.

**Figure 4 -.**
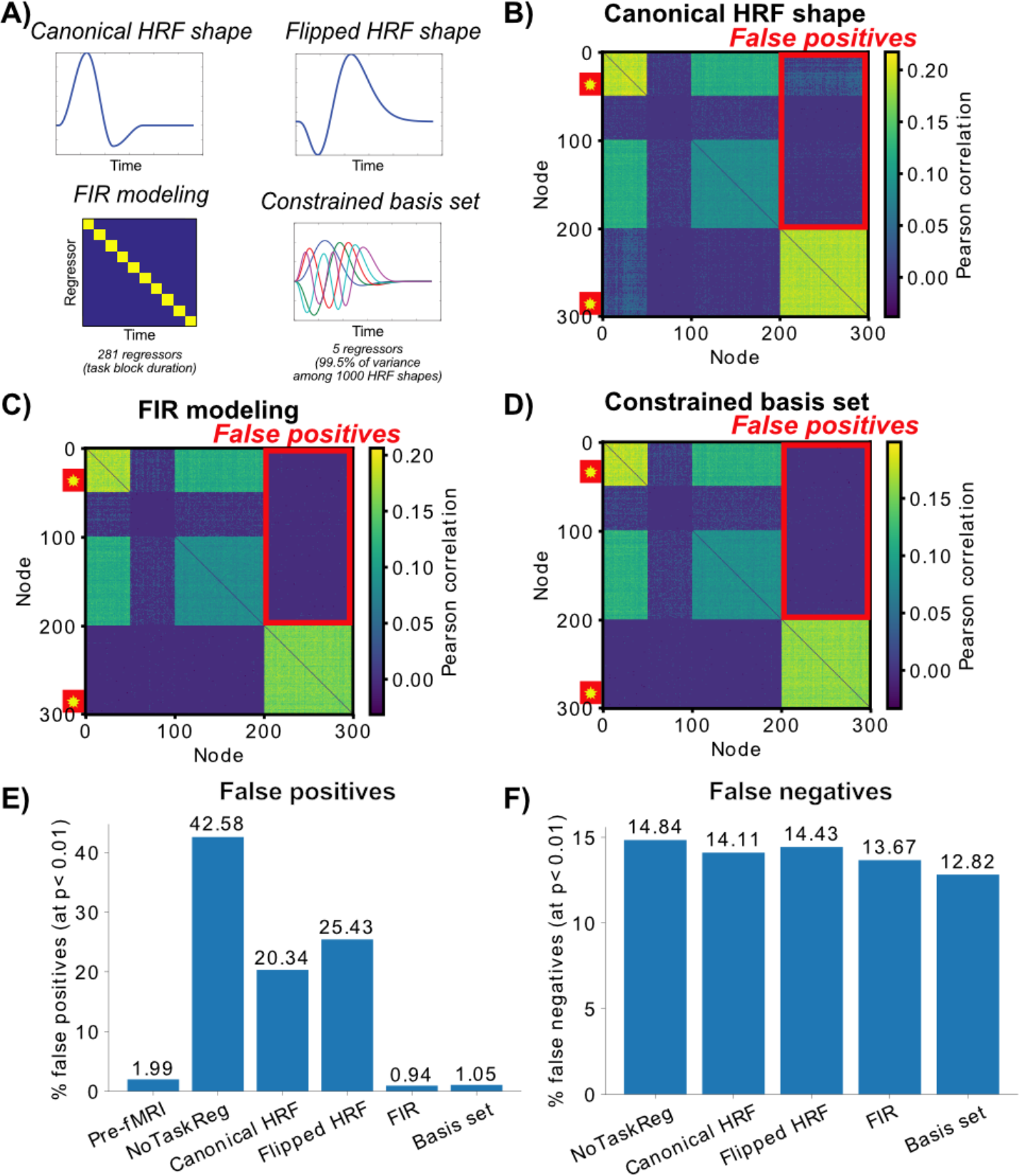
Testing task-timing regression approaches to reduce false positive rate. While some researchers investigating task FC fMRI ignore this problem, there are several standard approaches for attempting to reduce potential false positives. Critically, the 300-node computational model can provide a ground-truth scenario for testing the validity of these approaches. Note that all approaches are designed to leave moment-to-moment (and event-to-event) task-related variance in the time series, but to remove cross-event responses related to the task’s timing. Task vs. rest Pearson correlation differences (t-test p<0.01 thresholded) are shown. **A**) The 4 tested approaches are illustrated. The canonical HRF shape is what is typically used to reduce false positives in the literature, as with PPI. To assess whether the HRF shape mattered a “wrong” HRF was also used. The finite impulse response (FIR) and constrained basis set approaches are flexible, allowing them to fit the actual HRF shape. **B**) The canonical HRF shape task regression. There was a reduction from the no-regression condition (42.58%) but the remaining high false positive rate (20.34%) demonstrates that task regression with the canonical HRF is helpful but fails to correct the problem. Results were highly similar for the “flipped” HRF shape version (not shown). **C**) Task regression with the FIR approach eliminates the problem, with the false positive rate just below the expected detection rate of 1% (given our p<0.01 threshold). **D**) Task regression with a basis set of 5 regressors (accounting for 99.5% of the variance among 1000 plausible HRF shapes) was also successful in reducing the false positive rate (1.05%). **E**) False positive rates across six variants of the analyses. Since results were thresholded at p<0.01, any values above 1% can be considered false positives. **F**) False negative rates across five variants, with the pre-fMRI/neural variant treated as the “ground truth”. The entire 300 x 300 connectivity matrix was included in this analysis (rather than just the no connectivity zone). The fMRI simulation resulted in false negatives due to temporal smearing and downsampling, yet task regression reduced these false negatives.

A standard approach for empirically determining the correct HRF shape for task regression is finite impulse response (FIR) modeling (Cordova et al., 2016; Fair et al., 2007; Norman-Haignere et al., 2012). This involves including a binary regressor for every time point in the task event/block (**Figure 4A**) and using the residual time series, which is virtually identical to simply subtracting the mean evoked response (see **Figures 1C** & **2I**). This is sometimes referred to as “background connectivity” analysis when used with a block experimental design (as here) (Cordova et al., 2016; Norman-Haignere et al., 2012). As expected, we found that FIR modeling successfully reduced the false positive rate below the 1% specified by the p-value threshold (p<0.01): 0.94% false positive rate.

The success of the FIR approach suggested that flexibly fitting each region’s (for each subject’s) HRF shape was critical for correcting the false positive rate. We next tested this hypothesis more fully by using an alternative approach that also flexibly fits HRF shapes, but with fewer regressors. This approach - the constrained basis set approach (Woolrich et al., 2004) - involves reducing many plausible HRF shapes (variants on the canonical HRF) to a select set of basis functions using singular value decomposition. Note that the first three regressors included as basis functions were highly similar to the canonical, temporal derivative, and dispersion derivative regressors (respectively) commonly used with SPM software (Woolrich et al., 2004).

Consistent with our hypothesis, we found that the constrained basis set approach also reduced the false positive rate to the level expected with the p-value threshold (p<0.01) (**Figure 4E**): 1.05% false positive rate. These results confirm that flexibly fitting each region’s HRF was important for reducing the false positive rate, though it appeared that the basis set approach was somewhat less effective than the FIR approach.

### Neural mass model: Testing for potential false negatives due to false positive correction strategy

Given that the approaches involving more regression parameters did better, it is possible that the reduction in false positives was due to removing variance generally (rather than just the variance associated with false positives). This possibility predicts that the FIR and constrained basis set approaches would inflate false negatives along with reducing false positives. We began by setting a baseline by comparing no-task-regression fMRI FC estimates to no-task-regression neural FC estimates (i.e., the data prior to HRF convolution). This isolated the effect of the fMRI simulation on the FC results, given that fMRI simulation was the only difference between these two conditions. We found a 14.48% false negative rate (along with a 19.31% false positive rate) for no-task-regression fMRI FC relative to no-task-regression neural FC. Based on this, a 14.48% or lower false negative rate when using the FIR or basis set approach would indicate that these approaches did not increase the false negative rate (**Figure 4F**).

As expected, the false negative rate for the FIR and basis set approaches were both below 14.48%: 13.67% for FIR and 12.82% for basis set. These results suggest that the FIR and basis set approaches removed variance that was inappropriately altering FC estimates, both in terms of false positives and false negatives. Note that, when using the entire FC matrix (rather than just the no-connectivity zone), the false positive rate dropped from 19.31% for no-task-regression to 0.75% for FIR and 0.75% for basis set approaches - smaller false positive rates than observed when focusing solely on the no-connectivity zone. Together these results suggest that the extra regression parameters included in the FIR and basis set approaches are unlikely to reduce false positives by also reducing true effects (and that they can actually increase detection of true effects).

### Empirical fMRI data: Testing the efficacy of false-positive-reduction approaches

We next tested the ability of the FIR and constrained basis set approaches to reduce task-state FC false positives relative to other standard false-positive-reduction approaches (see **Table 1**). Unlike the computational models, we did not know the “ground truth” here, so we had to rely on any reduction in detected task-state FC as a proxy for false-positive reduction. Importantly, the FIR and constrained basis set approaches are unlikely to create false negatives given that they did not inflate the false negative rate in the neural mass model (**Figure 4F**).

A set of 360 functionally-defined nodes were used (Glasser et al., 2016) (**Figure 5A**) to calculate cortex-wide FC across seven highly distinct tasks in 100 healthy young adults. Without task regression the percentage of connections that increased from resting-state FC to task-evoked FC (false discovery rate corrected for multiple comparisons) was 7.22% across the seven tasks (**Figure 5B**). Only slightly reduced values were found for task regression with the canonical HRF approach (4.90%). Critically, there were substantial reductions in the percentage of task-state FC increases when using the FIR (2.49%) and constrained basis set (3.01%) approaches. These results suggest that the model results presented previously likely showed a “worst case” scenario, but that false positives can nonetheless almost triple the rate of detected task-state FC changes when an effective task regression approach is not used. See Supplementary Materials for more details.

**Figure 5 -.**
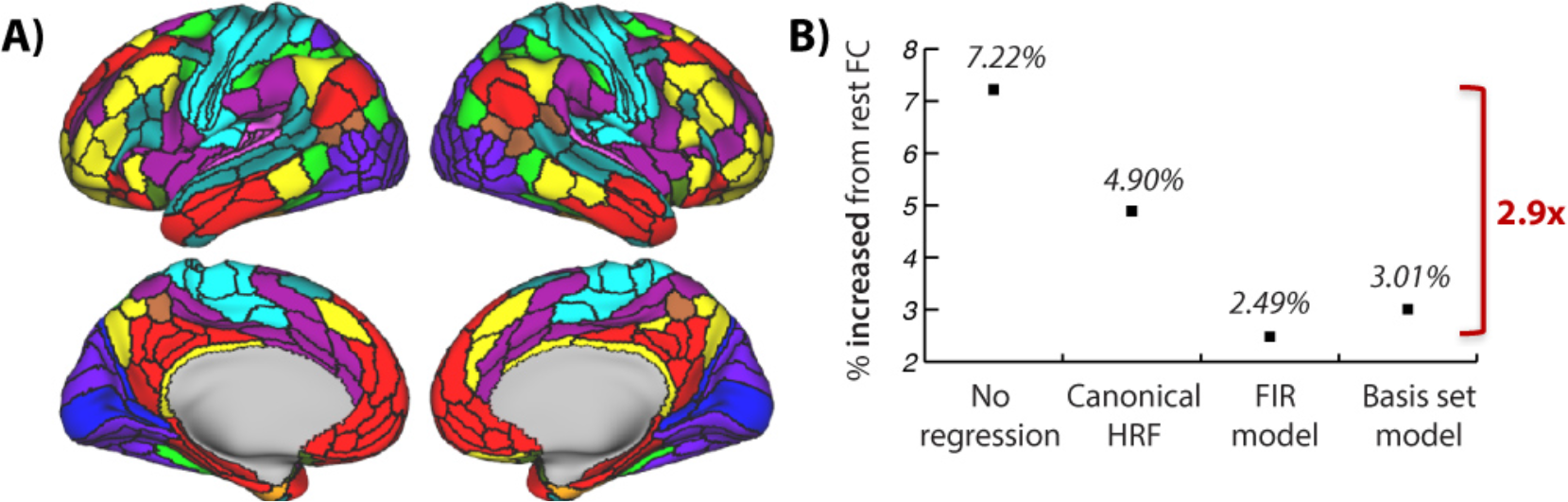
Analysis of empirical fMRI data reveals likely false positive rates for task-state FC estimates (with resting-state FC as a control for spontaneous correlations) **A**) The regions used for data analysis, as defined by Glasser et al. (2016). Colors reflect functional network assignments used for FC matrix visualization (Spronk et al., 2017) in subsequent figures. These assignments were used solely for visualization - results were not affected by the chosen network assignments. Colors match the network labels in Figure 6. **B**) The cross-7-task average rate of significant task-state FC increases from resting-state FC are shown (using Pearson correlation, FDR corrected for multiple comparisons, p<0.05). To the extent that the FIR approach eliminates false positives (demonstrated in the neural mass model), the percentages suggest a false positive rate of 65.5% without task-regression preprocessing, 49.2% with canonical HRF and 17.3% with constrained basis set model approaches. There were 2.9 times more significant FC increases without task regression compared to when FIR task regression was used. Note that resting-state FC is used here simply as a baseline (to control for FC driven by spontaneous activity) rather than as the ground truth FC.

Given that FIR modeling was most effective in reducing false positives in the neural mass model, and reduced task-state FC estimates the most in the empirical data, we identified FIR as the preferred method. This conclusion is further supported by a supplementary analysis found that FIR task regression reduced inter-subject correlations (time series correlations primarily driven by task timing) the most among the tested methods (**Figure S1**). Having identified FIR as the preferred method, we next quantified the amount of likely task FC inflation by comparing the no-task-regression task FC estimates versus task FC estimates with FIR-based task-timing regression. This involved comparing each connection’s FC value without task regression to its FC value with FIR regression. **Figure 6** plots these statistically significant (p<0.05, FDR corrected) differences for all seven tasks individually. The percentage of connections with significant (p<0.05, FDR corrected) differences for each task are reported in **Table 3**, with significant increases and decreases in FC strength between the approaches listed separately. Note that these false positive estimates were highly similar to false positive estimates obtained using a non-parametric shuffling procedure often used for correcting task-timing confounds in spike correlation studies (see Supplementary Materials, **Figure S2**). Additionally, we found substantial task-timing-induced FC inflation with PPI analysis relative to the FIR approach (see Supplementary Materials). These results demonstrate that task-timing regression matters in practice, as it significantly alters task-state FC estimates across a broad variety of brain regions across a broad variety of task manipulations.

**Table 3 -.** Amount of likely activation-induced task FC inflation. Comparison of no regression vs. FIR regression approaches, listed for each of the empirical fMRI tasks. The numbers indicate the percentage (out of all possible pairwise connections) of significant task FC differences (p<0.05 FDR corrected), comparing FC estimates between no-task-regression and FIR-task-regression approaches. These percentages are based on the values plotted in Figure 6, but separating increases from decreases in FC estimates with no task regression (relative to FIR regression). For example, 10.9% increased connections for the Emotion task indicates that 10.9% of all 64,620 connections were significantly larger between no task regression vs. with FIR regression. A similar table reporting PPI inflation can be found in the Supplementary Material (Table S1).

| Task name | % connections increased with no-task-regression | % connections decreased with no-task-regression |
| --- | --- | --- |
| Emotion | 10.9% | 7.2% |
| Gambling | 36.0% | 8.9% |
| Language | 27.9% | 25.0% |
| Motor | 29.7% | 3.6% |
| Social | 41.5% | 13.2% |
| Reasoning | 49.8% | 15.9% |
| Working memory | 28.3% | 8.8% |

**Figure 6 -.**
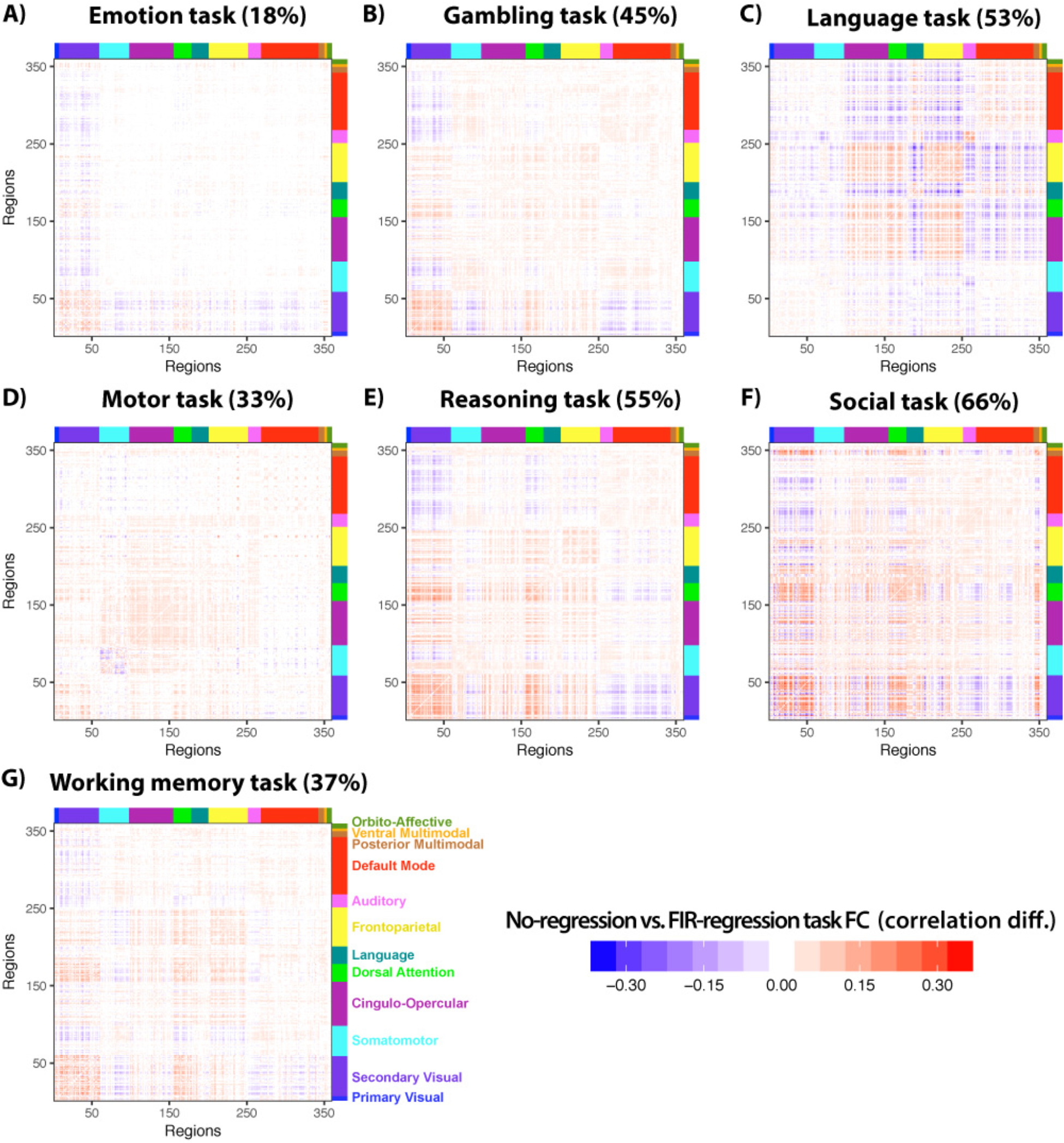
Estimated FC inflation for each of the 7 tasks. Task-evoked activation-based FC inflation was estimated by contrasting no-regression from FIR-regressed task FC estimates. Only statistically significant (p<0.05, FDR corrected) differences are shown for each task. Each FC matrix is shown with the name of each task and the percentage of connections (of the entire 360 x 360 FC matrix) that were significantly different between the no-regression and the FIR-regressed task FC estimates. Note that all tasks involved visual stimuli except for the language task.

### Empirical fMRI data: Testing for generalization to task-to-task FC changes

The prior results demonstrate inflation of task-state FC, suggesting that task-to-task FC differences would also be altered. This result was not guaranteed, however, given the possibility that the task-state FC inflations reported above were subtle and therefore only detectable for large cognitive contrasts (such as between task and rest). We tested for cross-task alterations in the well-studied N-back task’s 2-back vs. 0-back contrast (Barch et al., 2013). This is one of the seven tasks included in the prior analyses, now with the 2-back and 0-back conditions estimated separately.

As expected, we found that results were similar with the cross-task FC comparison as the task-to-rest FC comparison. Specifically, the approaches that flexibly modeled the HRF shape (FIR and basis set approaches) produced fewer significant results than alternate approaches (**Figure 7**). Without task regression the percentage of connections with task-state FC changes (false discovery rate corrected for multiple comparisons, p<0.05) was 28.14% (**Figure 7A**). Only slightly reduced values were found for task regression with the canonical HRF approach (24.97%; **Figure 7B**). Consistent with the task-to-rest FC comparison results, there were substantial reductions in the percentage of task-state FC increases when using the constrained basis set (12.92%; **Figure 7C**) and FIR (2.89%; **Figure 7D**) approaches. In contrast with the task-to-rest FC comparison results, however, FIR regression reduced the number of significant results relative to the basis set approach (2.89% vs. 12.92%). Notably, the significant reduction of visual network FC with the dorsal attention network (from 2-back to 0-back) was present for three of the methods but went away with FIR regression - the method that most flexibly fits HRF shape and thus likely best reduces false positives. This demonstrates a large-scale conclusion that could have been reached erroneously if FIR regression was not used to remove task activations.

**Figure 7 -.**
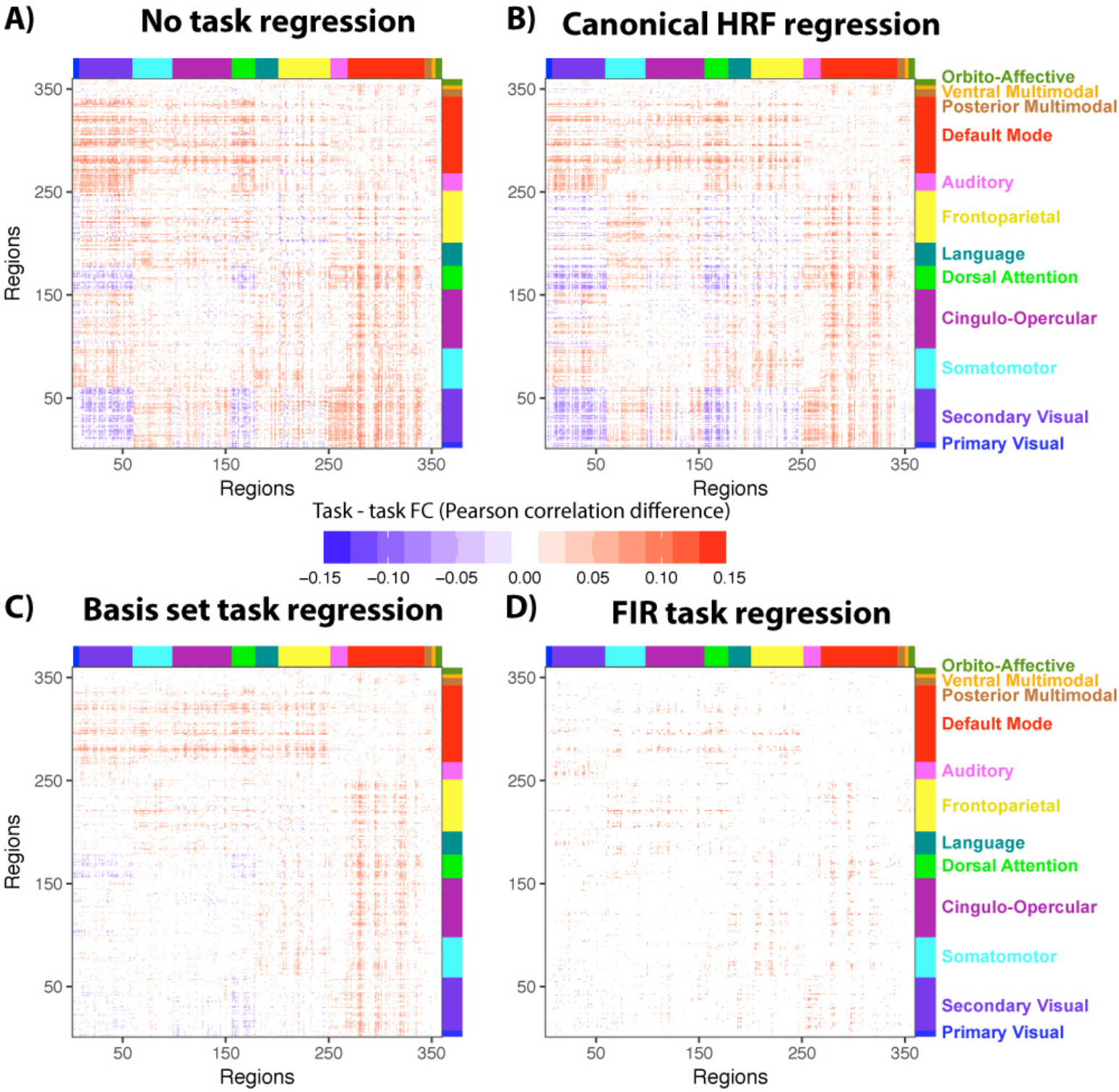
Task-to-task FC comparison: 2-back vs. 0-back (N-back working memory task) **A**) 2-back vs. 0-back FC differences, with no task regression preprocessing (p<0.05, FDR corrected for multiple comparisons). **B**) Identical to panel A, but with constrained basis set task regression preprocessing. **C**) Identical to panel A, but with canonical HRF task regression preprocessing. Note the visual similarity to the no-task-regression results. **D**) Identical to panel A, but with FIR task regression preprocessing.

These results suggest that the small FC differences between well-matched task conditions can be more sensitive than task-to-rest comparisons to the quality of GLM fit for the FC pattern that emerges. Based on the neural mass model results indicating that the fMRI data better reflect neural (i.e., input/LFP) data when using the FIR approach, and the additional flexibility of the FIR approach (without inflated false negatives) relative to the basis set approach, we interpret the FIR results as likely being more accurate than the other approaches. Note, however, that (regardless of regression method) concluding a true change in FC occurred - rather than a change in unshared variance (e.g., noise) - would require additional tests such as unscaled covariance (Cole et al., 2016b).

### Empirical fMRI data: Visualizing the relationship between task co-activation and task-state FC inflation

We next sought to visualize the correspondence between mean task-evoked responses (as estimated using GLM analysis) and task-state FC inflation to help further empirically establish its robustness. First, we calculated task-state FC inflation as the difference between no-regression task FC and FIR-regressed task FC. We then visualized this difference for all connections for an example task - the “working memory” HCP task (**Figure 8A**). The working memory task was chosen as the example task due to there being more data per subject for that task than the others (increasing statistical power). This revealed that much of the task-state FC inflation was related to visual network connections, consistent with this being a task involving visual stimuli. Notably, not all connection changes were positive, suggesting that co-activations in the opposite direction (e.g., a positive activation for one region and a negative activation in the other) could lead to artificial FC reductions. We verified that this is a likely explanation for FC reductions by visualizing the FC inflation results alongside the actual activation pattern (**Figure 8A**). Specifically, it appeared that negative activation in default-mode network regions (see upper portion of activation vector in Figure 8A) led to under-estimated FC with the positively-activated visual network regions (see blue values in upper-left of the Figure 8A FC matrix).

**Figure 8 -.**
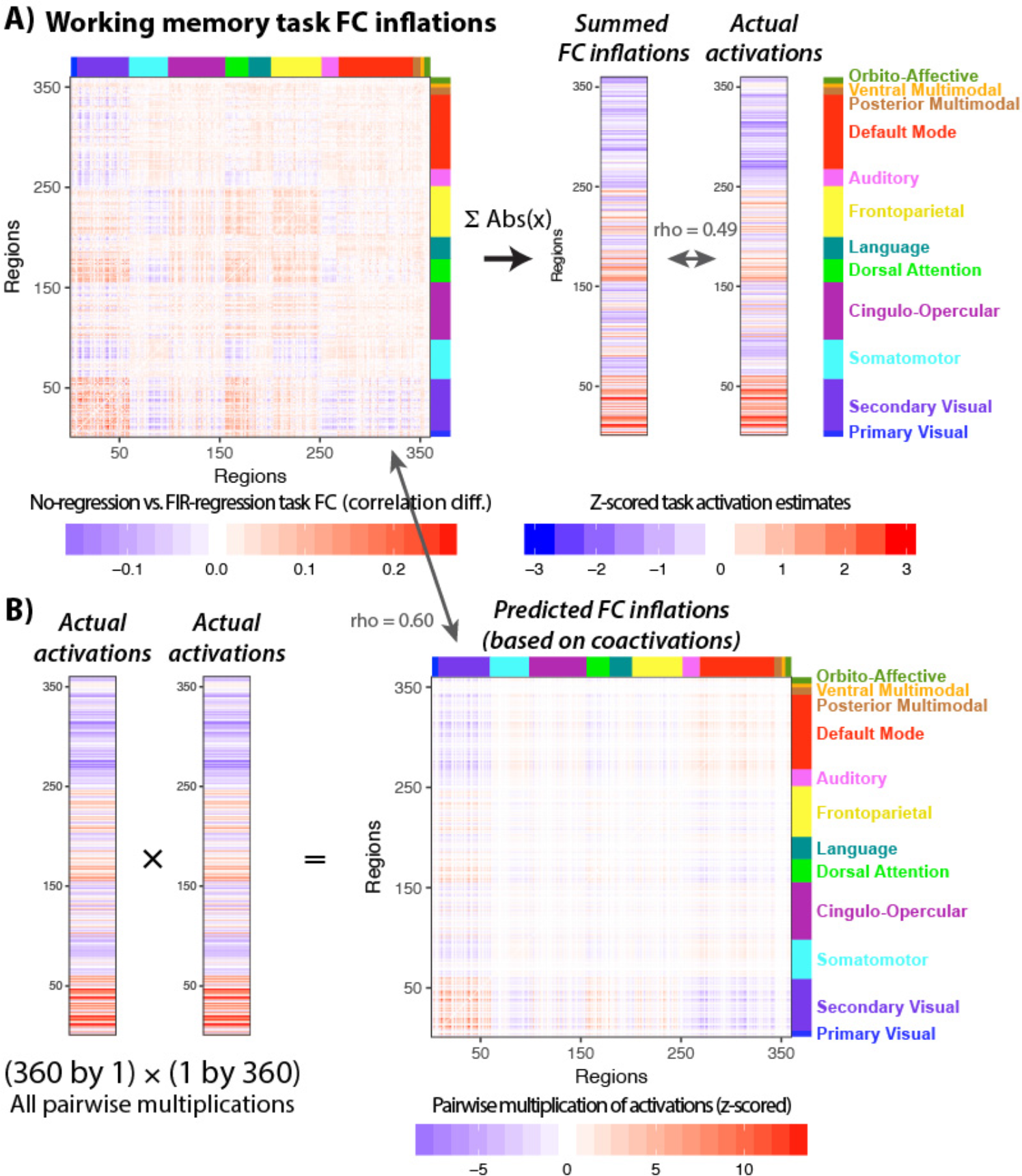
Visualizing the relationship between task co-activation and task FC inflation. **A**) Task-state FC inflation is shown (left) by subtracting the group-mean FIR-regressed task FC matrix from the group-mean non-regressed task FC matrix. An example task - the HCP “Working memory” task (which involves visual stimuli and button pressing) - is used for illustration (with no thresholding). The FC inflation values were summed (after taking the absolute value) by region to summarize the degree to which each region showed FC inflation. This was then compared with the task-evoked activation pattern (estimated using a standard GLM with a canonical HRF shape), showing a significant correspondence (Spearman rank rho=0.49, p<0.0001). This provides a way to visualize the degree to which co-activation patterns are likely influencing task FC patterns. **B**) The group-mean task activation pattern was used to predict likely inflation of task-state FC estimates driven by co-activations. This involved multiplying each activation with all others in a pairwise manner, converting the activation vector into a co-activation matrix. There was a significant similarity between the co-activation matrix and the task-state FC inflation (Spearman rank rho = 0.60, p<0.0001). This shows an alternative way to visualize the degree to which co-activation patterns are likely influencing task FC patterns.

We next sought to create a simple summary of the task-state FC inflation by region, so it could be compared directly to the task activation pattern. This involved summing the task-state inflation values by region (i.e., summing across all the columns in the task-state FC inflation matrix for each row), after taking the absolute value for each number. This is visualized for the example task in **Figure 8A** (see right ‘Summed FC inflations’ vector). We found that this simple summary correlated significantly with the actual activation pattern for all seven tasks, all p<0.00001 except for Task 3 (the “language” task; p=0.0003). The Spearman rank correlation rho values for each task, respectively: 0.35, 0.31, 0.19, 0.47, 0.56, 0.56, 0.49. These results demonstrate the robustness of the association between task-evoked activations and task-state FC inflation.

To further illustrate the relationship between activation and task FC inflation, we next sought to create a simple prediction of task-state FC inflation based only on the observed co-activation pattern. Task-state co-activation inflation was conceptualized simply as the pairwise product of the task GLM estimates. Multiplying the activation values results in cases wherein large positive co-activations are expected to create the largest increases in task FC estimates. In contrast, co-activations in the opposite direction (e.g., a positive activation and a negative activation) are expected to cause task FC estimate decreases. This sort of prediction is visualized for the example task in **Figure 8B**, showing robust correspondence with the actual task-state FC inflation pattern (**Figure 8A**). This correspondence between the predicted and actual task-state FC inflation was statistically significant across all seven tasks (all p<0.00001). The Spearman rank correlation rho values for each task was, respectively: 0.51, 0.74, 0.04, 0.28, 0.67, 0.60, 0.60. Note that the third (“language”) task was still statistically significant despite having a small effect size, given the large N when comparing entire FC matrices (64,620). These results further demonstrate the robust association between task-evoked activations and task-state FC inflation, this time by starting from the co-activation patterns to show how even complex patterns of FC can be driven by activation-based inflation. Note that we did not expect exact correspondence between the predicted and actual task-state FC inflations, given that (among other factors influencing FC inflation) HRF shape is known to vary across regions, which likely adds noise and reduces FC inflation.

## DISCUSSION

We found strong evidence that task-evoked activations led to spurious but systematic changes in fMRI-based task FC estimates. This was noted as a possibility in previous publications (Al-Aidroos et al., 2012; Cole et al., 2013; Fair et al., 2007; Friston et al., 1997; Gratton et al., 2016) but, to our knowledge, has never been conclusively established theoretically (using computational modeling) or empirically. Further, this hypothesized issue with task FC has typically been described generally, without reference to it being particularly problematic for fMRI analyses. We began by modeling the hypothesized effect using two computational models. Notably, we did not force the models to show activation-induced FC inflation, but discovered that it emerged simply from modeling fMRI task activations. Critically, beyond merely demonstrating the extent of the task-state FC inflation, we also evaluated the efficacy of different correction methods. Regression methods that flexibly fit hemodynamic response shape - FIR and basis set GLM approaches - were found to eliminate activation-induced FC inflation (without increasing false negatives), whereas alternative methods did not. Consistent with these theoretical results, we found that FIR and basis set approaches significantly reduced task FC estimates in empirical fMRI data. We found that the FIR approach reduced task FC estimates the most, consistent with its unique ability to flexibly fit any possible HRF shape, suggesting this as the preferred approach.

**Table 4 -.** Frequently asked questions. The variety of considerations relevant to the topic of mean task-evoked responses potentially inflating task-state FC estimates are substantial. We therefore include this question-driven organization of relevant considerations to provide a more direct means for frequent questions to be addressed.

| Questions | Answers |
| --- | --- |
| 1) Why remove mean evoked responses prior to task-state FC analysis? | So that the amount of inter-node linear interaction can be estimated, above and beyond mere task co-activations. Rather than the ambiguous inference that two regions are "likely active or interacting during the task", removing mean evoked responses allows for the more specific inference, "likely interacting during task". Given that most neuroscience studies have focused on mean evoked responses, it is important to remove these responses to reduce the risk of reporting mean evoked responses (mere first-order changes in activity level) as "connectivity" results (second-order effects). |
| 2) What other reasons are there for removing the mean evoked responses prior to task-state FC analysis? | Task-state FC can be conceptualized as an interaction between psychological and physiological factors – a "psychophysiological interaction" (PPI) (Friston et al., 1997; McLaren et al., 2012). As with most interactions, it is important to account for the main effects prior to interpreting the interaction. This ensures the estimated interaction is not just the main effect (here, mean evoked response) masquerading as an interaction. Additionally, task-state FC can be conceptualized in terms of causal inference, in which the main effect of task events is a confound for estimating the interaction between pairs of nodes. This is clearly demonstrated in the "no connectivity zone" example in Figure 3, wherein the main effect produces correlations in nodes with no physical means of interacting. |
| 3) How can the task-timing confounding be corrected? | As with all confounds, holding the confounding factor constant can remove its effect (Pearl, 2009). The most straightforward strategy for holding stimulus/task state constant is to only include time points from that state – accounting for the main effect of task state. Critically, however, neural processing involves temporal autocorrelation, such that the effects of rest-to-task (or task-to-task) state changes extend beyond stimulus presentation (state-transition transients). Both models reported here indicate that the temporal autocorrelation introduced by the fMRI BOLD response is especially problematic in this respect (see Figure 2). Another strategy is therefore necessary, and we advocate subtraction of an estimate of the temporally-extended response to the stimulus. We also show that flexible fitting of the task-evoked response is critical for properly removing the task-timing confound for fMRI analyses. |
| 4) Does removing the mean evoked response remove too little variance (leaving false positives)? | This is unlikely, given that all false positives were removed in the minimal and neural mass models following subtraction of mean evoked responses. This result appears to rely on several prerequisites: 1) stimulus identity and timing are identical across event instances, 2) enough time points are included in the mean evoked response estimate to cover the temporally-extended response to the stimuli, 3) more than one event instance is included in the mean evoked response estimate (to allow separation of each evoked response from the mean response), 4) an accurate estimate of the evoked response shape is used (e.g., using FIR regression), and 5) enough temporal separation between events of different conditions to ensure separate mean evoked response estimates and correlation |
|  | estimates. Temporal jitter between stimuli can be useful for mean evoked response estimation (especially when linear regression is used) (Miezin et al., 2000), but this is not helpful for correlation estimation. Block designs (with or without temporal jitter) may be most appropriate, given that this allows removal of both transient and sustained evoked responses (Al-Aidroos et al., 2012; Visscher et al., 2003) with enough temporal separation between condition types. |
| 5) Does removing the mean evoked response remove too much variance (inducing false negatives)? | This is unlikely, given that the neural mass computational model indicated a *reduction* in false negatives following removal of the mean evoked responses. Yet it is technically possible that two nodes could have a highly stereotyped interaction time-locked to the stimuli, producing covariance that is included in the mean evoked response. In this case, removing the mean evoked response from both nodes would inappropriately reduce the detected covariance. However, since neural populations are known to always have variability across event instances, moment-to-moment and event-to-event covariance would remain, such that the proper level of covariance would likely be detected. Supporting this conclusion, we found that ~90% of inter-region covariance remained in fMRI data after removing the mean evoked response from all nodes (see Supplementary Materials). In other words, if two nodes are truly interacting, evidence for such interaction is very likely to be present in the moment-to-moment and event-to-event variance left over after removal of the mean evoked responses. |
| 6) Do other sources of confounding remain after removal of mean evoked responses? | Yes. Any time a node influences two or more other nodes there is likely an inflation of the correlation between those downstream nodes. While this is problematic, we focus here on the confounding influence of external stimuli. It will be essential for future research to resolve the more general issue of confounding in FC analysis. Some progress toward removing such confounds have been made, typically via partial correlation or multiple regression (Cole et al., 2016a; Ramsey et al., 2010; Smith, 2012). Note that partial correlation and multiple regression FC approaches are likely only minimally impacted by the task-timing confound, due to partialling out of common signals across time series (mean evoked responses). |
| 7) Is it problematic that removing the mean evoked response leaves some evoked variance? | No. Subtracting the mean evoked response removes the first-order (main) effect of the task stimuli, leaving second-order event-to-event variability in the evoked response that can be used to assess state-dependent changes in the statistical relationship between nodes (i.e., task-state FC). Testing for correlations using such event-to-event variability is very similar to the "beta series" task-state FC approach, which estimates correlations between event-to-event variation in estimated evoked responses (Rissman et al., 2004a). |

### Why event-averaged task activation variance should be removed prior to estimating task FC

Our extensive computational and empirical investigation of activation-induced FC inflation suggests several reasons why event-averaged task activations should be removed prior to estimating task FC. For instance, we found that FC changes and activation amplitude changes are statistically and mechanistically distinct, such that they have meaningfully distinct implications for neuroscientific theory. Specifically, event-averaged task-evoked activations involve consistent cross-event activity amplitudes, while task-state FC involves synchronous moment-to-moment changes in activity indicative of direct or indirect neural interactions. This distinction can also be thought of as task-evoked activation being enhanced by low variance (amplitude consistency) contrasting with task-state FC potentially being enhanced by high variance (moment-to-moment covariance). Thus, even if one finds event-averaged task-evoked activation patterns of interest, they should be investigated separately from task-state FC due to this mechanistic distinction between them. Indeed, there are already sub-fields investigating task-evoked activation patterns - multivariate pattern analysis (Norman et al., 2006) and standard GLM analysis (Poline and Brett, 2012) - again supporting the conclusion that such effects should be isolated from task-state FC estimates.

Another reason to remove task activations prior to estimating task FC is that allowing task-evoked activations to inflate task-state FC estimates leaves open the possibility that new task-state FC effects are simply relabeling previously-discovered task-evoked activation effects as “connectivity”. This suggests that some previously-discovered effects that either did not remove any task activation variance, or that used a suboptimal approach for removing task activation variance, could have been driven to some extent by task-evoked activation changes. Notably, a handful of studies have already used FIR GLM to remove task activation variance prior to estimating task FC (Al-Aidroos et al., 2012; Cordova et al., 2016; Fair et al., 2007; Gratton et al., 2016; Norman-Haignere et al., 2012; Sadaghiani et al., 2015; Summerfield et al., 2006), suggesting these studies did not suffer from the task FC inflation effect identified here. Some have labeled this FIR-based removal of task activation variance followed by task FC estimation “background connectivity” (Al-Aidroos et al., 2012; Cordova et al., 2016; Griffis et al., 2015a; Norman-Haignere et al., 2012). The present results suggest “background connectivity” and related approaches are effective in reducing (and likely even eliminating) task FC false positives driven by fMRI task activations.

A skeptic might argue that one could reverse this argument, with task FC being the real effect and task activations being secondary. The computational model analyses here demonstrate this is incorrect, since there are cases in which no true task FC is possible yet task FC is spuriously detected due to task co-activation (see the “no connectivity zone” in Figure 3). Further, it is clear that task activation is the first-order effect (simple change in cross-event mean amplitude), whereas task FC is a second-order effect building on covariance in moment-to-moment activation amplitudes. It is customary in science and statistics to account for simpler, first-order effects prior to interpreting second-order effects, such as interpreting ANOVA interactions only after accounting for main effects. Thus, an effect that can be explained as either a task activation or a task FC change would be preferentially interpreted as the simpler of the two - a task activation.

Another concern of a skeptic might be that removing task activation variance would remove the very task FC effects s/he is interested in. Both the model and the empirical results demonstrate that this is highly unlikely. First, we found that FIR task regression did not increase the rate of false negatives in the neural mass model. Indeed, FIR task regression *reduced* the rate of false negatives, suggesting FIR task regression might even increase the number of detected true task FC effects (rather than simply reducing false positives) (see **Figure 4F**). Second, we found that the event-averaged task activation variance removed was only a small percentage (~10%) of the shared variance in the empirical fMRI data (see Supplementary Materials), suggesting that the bulk of the effects without activation regression was already driven by moment-to-moment variance independent from event-averaged activations. This suggests that even those who interpreted task FC in terms of event-averaged co-activation were actually observing primarily correlations of moment-to-moment fluctuations. Notably, despite most of the variance being driven by moment-to-moment fluctuations, we found that event-averaged activations alter task FC estimates substantially enough that many false conclusions are obtained without first removing event-averaged task activation variance. These two findings - that task activations were both a small portion of the overall variance and made a meaningful difference to results - can be reconciled by considering that a relatively small percentage of false positives among thousands of functional connections would nonetheless produce a large number of false inferences.

Taken together, these considerations suggest removal of mean task activation variance from neural time series does not remove the covariance of interest when investigating task-state FC. Yet, what might the covariance of interest represent, mechanistically? There are several possibilities. First, it is likely that spontaneous covariance present during resting state is also present during task performance. Evidence for this comes from many sources, such as the observation that neural signals are more dominated by spontaneous than task-evoked activity during task performance (Raichle:2010bt; Raichle and Mintun, 2006), and the observation that the spontaneous activity during task performance has a similar correlation structure as during resting state (Cole et al., 2014; Fox et al., 2007; 2006; Krienen et al., 2014). This source of covariance is not expected to differ much (if at all) from resting state, which is why we chose to use resting state as a control condition for several analyses to better isolate task-state-specific FC effects. Second, another likely source of task-state covariance is extended state-related changes in neural interactions, such as from activity-induced short-term (or long-term) plasticity of synapses (Karmarkar and Dan, 2006) or sustained activity in one area influencing others (Miller and J. D. Cohen, 2001). Finally, any neural processes that vary in their timing and/or amplitude across events will remain after cross-event mean evoked activity regression. Since virtually all neural processes vary in their exact timing and/or amplitude across events (due to the stochastic nature of neural activity), nearly every neural process will ultimately be included (potentially with some attenuation) after cross-event mean evoked activity regression. Among these neural processes, those that result in non-zero covariance with other measured neural processes (such as via long-distance neural interactions) will result in a non-zero task-state FC estimate.

### Limitations and opportunities for further research

As with most studies, many possible analyses related to the core research question were not included here, providing opportunities for future research. For instance, it could be informative to use a neuron-level computational model to further verify the results obtained using the neural mass model (Brette et al., 2007; Goodman, 2008). However, our neural mass model was intentionally kept simple and abstract, with the expectation that this abstraction will increase the probability that results will generalize to many different possible computational models (including highly realistic neuron-level models). The key idea is that abstraction to neuron-like units in the neural mass model reduces the number of assumptions by identifying key effects that are general enough to emerge from properties present in a variety of neuron-like interactions (e.g., across spatial scales). Despite the plausibility of this expectation it is of course important to test this prediction using more detailed neuron-level modeling.

There were several aspects of the computational model results that did not completely agree with the empirical fMRI results. First, we empirically observed more task-state FC decreases from resting state, whereas the computational model results showed more task-state FC increases from resting state. This likely reflects our use of task-state FC increases from resting state (among the first 50 nodes) to select the model parameters. Notably, in the model we saw task-state FC decreases between the first 50 and second 50 nodes, due to there being inhibitory connections between those two network communities. This could suggest that more inhibitory connectivity should have been included in the model in order to match the empirical results. Alternatively, we could have selected model parameters based on maximal decreases in task-state FC relative to resting state. This may have resulted, for instance, in a higher bias parameter, equivalent to a larger amount of spontaneous activity leading to larger resting-state correlations. This issue is related to improving understanding of why the empirical results showed that most functional connections are lower during task relative to rest.

Another aspect of the computational model results that was not in complete agreement with the empirical fMRI results was the observation that FIR task regression reduced task-state FC estimates substantially more than basis set task regression. In the model the basis set approach involved only 1.05% false positives, very similar to the 0.94% false positives with the FIR approach. While the results were similar for task-state FC vs. resting-state FC (2.49% detected effects with FIR vs. 3.01% with basis set), our task-to-task FC comparison indicated substantially fewer detected FC differences when using FIR (FIR: 2.89%, basis set: 12.92%). Given the more flexible fitting of HRF shape with FIR, it is likely that FIR task regression better fit and removed the task-evoked activations than the basis set approach. It is possible that the extra flexibility of FIR over fit the task-evoked time series, removing additional noise but also some covariance of interest. However, the computational model results suggest that, if anything, this extra flexibility likely reduced (rather than increased) false negatives, potentially by removing more noise than signals reflecting true interactions. It will nonetheless be important for future research to quantify the degree to which FIR model overfitting results in inflation of false negatives in empirical results.

We were able to use the computational model to conclusively show that coactivations can induce spurious fMRI task FC by creating a “no connectivity zone” wherein no true task FC can be possible. Ideally, however, we would have had this sort of scenario in the empirical fMRI dataset. Instead, the empirical fMRI analyses supported the plausibility of task FC being inflated, with detected increases and decreases in task FC once event-averaged task activation variance was removed. This leaves open the possibility (however small) that removing cross-event mean responses removed some true task Fc effects. It will be important for future studies to investigate this possibility. Notably, however, the computational modeling results demonstrated that false negatives were not increased (and were in fact decreased) when cross-event mean responses were removed. Again, this suggests that, if anything, removing cross-event mean responses in turn increases the number of true task FC effects detected (rather than decreasing them).

We focused primarily on Pearson correlation-based task FC. It will be important for future research to test the generality of our conclusions to all task FC approaches. We showed that the results at least generalize to PPI analyses, suggesting the findings will likely generalize further. Indeed, the generalization to PPI suggests the task FC inflation effect is driven primarily by a change in covariance - the quantity underlying a variety of association measures used for task FC analysis (such as Pearson correlation and PPI) (Cole et al., 2016b). This is consistent with the minimal model results (Figure 2), which shows that the underlying task FC inflation is driven primarily by similarity in the hemodynamic response. Such clear similarity - which was induced by convolution with a similar-shaped HRF - suggests this effect will generalize such that a variety of task FC measures will be inflated by fMRI task co-activation.

It will be important for future research to investigate alternative approaches to correcting the task FC inflation seen here. For instance, one promising approach is blind deconvolution (Havlicek et al., 2011), which flexibly removes HRF shape from entire time series. This could, in theory, correct the inflation by estimating the true neural time series separated from the HRF. Such a result would be consistent with our finding that task FC was only minimally inflated in the neural time series in the computational model results. Another method that we expect to be effective in reducing or eliminating task activation-based inflation of fMRI task FC is the “beta series” task FC approach (Rissman et al., 2004a). In this approach, a separate GLM parameter estimate is fit to each task event (with an assumed HRF shape), with Pearson correlation of the parameter estimates (across voxels or regions) estimating task FC. In theory, this approach estimates task FC based on event-to-event (e.g., trial-to-trial or block-to-block) covariance (see Figure 1C), excluding most of the moment-to-moment covariance that is typically used. This approach’s use of an assumed HRF shape may result in false negatives (due to poor fit to activations in some cases), but appears unlikely to suffer from the same task FC false positives characterized here, given that beta series correlations isolate variation in evoked response amplitudes from evoked response shape. This suggests that studies that used beta series correlations are unlikely to have been influenced by the false positives characterized here (Cisler et al., 2014; Gazzaley et al., 2004; for example: Nee and Brown, 2012; Rissman et al., 2004b; Zanto et al., 2011), though future research will be important for verifying this. With regard to false negatives, Al-Aidroos et al. (2012) utilized a data-driven approach to identifying the shape of the evoked response, isolating event-to-event covariance without assuming a response shape. This might be an effective approach to improve estimation of event-to-event variance, possibly reducing false negatives in beta series analyses. Notably, however, Al-Aidroos et al. (2012) found that moment-to-moment covariance drove task-state FC estimates much more strongly than event-to-event covariance. Consistent with a minimal role for event-to-event variance, Fox et al. (2007; 2006) found that event-to-event variance is primarily driven by spontaneous moment-to-moment variance.

It will also be important for future research to investigate why the neural simulation (prior to HRF convolution) had some inflated task-state FC estimates. The inflation was quite small (a 1.99% false positive rate with a p<0.01 threshold), especially relative to the no-regression fMRI results (42.58% false positive rate), but it was nonetheless higher than expected by chance (1%, given the p<0.01 threshold). This likely reflects the small amount of coincident timing induced by the simultaneous stimulation across neural units, suggesting regression-based removal of task-evoked non-fMRI data (Headley and Weinberger, 2013; Karamzadeh et al., 2010; Mill et al., 2017) could also be useful for reducing false positives. Supporting this possibility, investigations of task-state FC with multi-unit recording in animal models (i.e., not involving the BOLD signal) have tended to remove cross-event mean evoked responses prior to estimating correlations among neural time series (termed “noise correlations”) in the interest of reducing false positives (Cafaro and Rieke, 2010; M. R. Cohen and Kohn, 2011). Demonstrating the equivalence of this issue for fMRI and non-fMRI data, we utilized a non-parametric approach based on methods popular with spike count correlations - involving shuffling events to estimate the contribution of confounding stimulus-evoked covariance (Averbeck et al., 2006; Grün, 2009) - with fMRI task-state FC estimation. These considerations make it clear that task-timing-induced correlation inflation is likely a problem for all forms of (direct or indirect) neural recording.

One remaining issue for the FIR GLM regression approach is that it relies on the particular set of regressors specified, when there might be additional task events unaccounted for. For instance, block onset and offset events with prominent fMRI activation responses have been identified (Dosenbach et al., 2006; Fox et al., 2005; Griffis et al., 2015b; Visscher et al., 2003), such that a standard FIR model of an event-related task design would fail to remove fMRI activation variance from these prominent events. The variance from these events would likely inflate task FC estimates. One solution would be to model these block onset and offset events separately so as to remove this variance prior to task FC estimation, as has been done recently (Griffis et al., 2015a). Another solution that was successfully applied here is to design task blocks of a given condition to have identical trial timings, then model all blocks with a single long set of regressors (such that all consistent within-block events would be modeled, including block onset and offsets) (Al-Aidroos et al., 2012).

Similar issues arise from rare events with large fMRI activation responses such as error trials (Menon et al., 2001; Neta et al., 2015) or learning-induced changes in activations, which are typically not accounted for separately in GLM models. Such events might also inflate task FC estimates, though they could also be included in an FIR GLM to reduce this effect. It will be important for future studies to consider these various scenarios and determine whether they can meaningfully alter task FC estimates. Given that most of the task FC inflation effect is caused by the HRF shape, another possibility would be to utilize blind deconvolution (Havlicek et al., 2011) to reduce this effect no matter its source (even those unknown to the experimenter). Another possibility is that the task-activation false positives arise solely from the experimental manipulation (task timing) acting as a confounding third variable, implying that internally-generated activation events (such as error trials or learning-related activation changes) reflect the brain dynamics of interest and therefore do not need to be removed. Notably, however, it seems likely that a region going from high activity to baseline-level activity with learning/practice would result in early activity having the activation confound but not later activity (resulting in learning-related task FC false positives when contrasting early vs. late activity). It will be important for future studies to investigate this issue, given the ambiguity (regarding false positives) of situations like error trials and task learning being an interaction between experimenter-induced task timing and internal processes.

A related issue is whether continuous task performance - such as continuous object tracking - eliminates the task-evoked activation confound for task-state FC estimation. Several task-state FC studies have utilized continuous task performance (Krienen et al., 2014; Tomasi et al., 2014), potentially to avoid the rest-to-task state transitions likely driving many of the false positives in the present study. However, it remains unclear whether events occurring during the continuous task constitute a task-timing confound, given that task-evoked activity would occur in response to these events. For instance, Tomasi et al. (2014) reported a continuous object tracking task in which two (of 10) objects were highlighted every 11.5 s. Subjects were instructed to press a button if the two highlighted objects were the tracked objects. Rather than being fully continuous, it is clear that these events would produce evoked activations in multiple brain regions (e.g., visual, motor, and somatosensory cortices), very likely creating task-timing-induced inflation of task-state FC estimates. Nonetheless, it is possible that such non-continuous events embedded within a continuous task would produce less of an FC inflation than, e.g., rest-to-task state transitions. It will be important for future studies to explore this possibility and, more generally, assess the promise of continuous task performance for reducing task-timing FC inflation.

### Conclusion

We identified strong evidence that fMRI-based (and to a lesser extent, non-fMRI-based) task FC estimates are consistently and spuriously altered by task activations. This was shown across a minimal model, a more realistic neural mass computational model, and empirical fMRI data involving seven highly distinct tasks. The models and empirical fMRI data analyses converged in suggesting that methods that remove event-averaged task activation variance - when flexibly taking HRF shape into account (especially FIR GLM) - are able to correct for activation-induced task FC inflation. These results suggest prior task FC fMRI studies that did not use FIR GLM as a preprocessing step might contain false positives. It will therefore be important to reanalyze data when possible, and begin using FIR GLM as a preprocessing step for task FC analyses moving forward. It might be tempting to retain event-averaged task activation variance in future task FC analyses given that the issue is not as problematic for non-fMRI data. However, the observation of inflated false positives in the “no connectivity zone” (1.99% with p< 0.01) for the neural non-fMRI simulation data suggests this is a fundamental problem for task FC analysis, such that task activation regression should be used with non-fMRI data as well. Moving forward, it will be important to develop a deeper understanding of why event-averaged task activation causes false positives even for non-fMRI data, as well as identifying alternative approaches to removing event-averaged task activations in both fMRI and non-fMRI data.

## Acknowledgements

The authors acknowledge support by the US National Institutes of Health under awards R01 AG055556 and R01 MH109520. Data were provided, in part, by the Human Connectome Project, WU-Minn Consortium (Principal Investigators: D. Van Essen and K. Ugurbil; 1U54MH091657) funded by the 16 NIH Institutes and Centers that support the NIH Blueprint for Neuroscience Research; and by the McDonnell Center for Systems Neuroscience at Washington University. The content is solely the responsibility of the authors and does not necessarily represent the official views of any of the funding agencies.

